# Dorsal-caudal and ventral hippocampus target different cell populations in the medial frontal cortex in rodents

**DOI:** 10.1101/2024.10.20.619325

**Authors:** Paola Alemán-Andrade, Menno P. Witter, Ken-Ichiro Tsutsui, Shinya Ohara

## Abstract

Direct projections from the ventral hippocampus (vHPC) to the medial frontal cortex (MFC) play crucial roles in memory and emotional regulation. Using anterograde transsynaptic tracing and *in vitro* electrophysiology in mice, we document a previously unexplored pathway that parallels the established vHPC-MFC connectivity. This pathway connects the dorsal-caudal hippocampus (dcHPC) to specific subregions of the ventral MFC, in particular the dorsal peduncular cortex. Notably, this pathway exerts a strong inhibitory influence on ventral MFC by targeting a substantial proportion of inhibitory neurons. Retrograde transsynaptic tracing in rats indicated that ventral MFC subregions project disynaptically back to vHPC. These results, altogether, suggest the existence of a remarkable functional circuit connecting distinct functional areas: the cognition-related dcHPC with the emotion-related ventral MFC and vHPC. These findings further provide valuable insights in the cognitive and emotional abnormalities associated with the HPC-MFC connectivity in neurological and psychiatric disorders.

## INTRODUCTION

Research over the years has shown that interactions between the hippocampus (HPC) and the frontal cortex play a crucial role in learning and memory processes. Overwhelming evidence suggests that abnormal connectivity between HPC and the frontal cortex is implicated in patients with neurological and psychiatric disorders who display cognitive deficits or disrupted emotional control^1,2^. This thus emphasizes the importance of more detailed investigations of the organization of the connectivity between these two cortical domains.

Recent functional neuroimaging studies suggest that the human HPC is connected with the frontal lobe and that this connectivity presents a functional topographical organization along the anterior-posterior axis of HPC^3–5^. Consistent with these observations are reports in non-human primates that HPC and the frontal cortex are unidirectionally connected by a direct projection that originates from field CA1 and subiculum (SUB) of HPC to the frontal cortex^6^. Although there is no consensus on which rodent brain regions correspond to the primate frontal cortex^7,8^, it has been shown that the medial frontal cortex (MFC), referred as medial prefrontal cortex in most rodent studies, receives massive inputs from HPC^9,10^. This rodent MFC can be functionally differentiated along the dorsoventral axis^11^. Whereas the dorsal MFC subregions, the dorsal anterior cingulate-(ACd) and prelimbic-(PL) cortices, play important roles in attention and working memory^12^, the ventral MFC subregions, the infralimbic-(IL) and dorsal peduncular-(DP) cortices, have been associated with emotional, autonomic and sympathetic functions^13,14^. Similarly, the rodent HPC has been functionally differentiated along its dorsoventral axis. Behavioral studies revealed that dorsal HPC plays a crucial role in spatial cognition whereas ventral HPC is involved in emotional-related behavior^15–17^ (See also Strange et al.^18^).

Given that rodent anatomical reports showed that the direct HPC-MFC projection originates largely from ventral HPC^9,10^, numerous behavioral studies have focused their attention to the ventral HPC projection to MFC. Indeed, the ventral HPC-MFC circuit is now thought to be involved in spatial working memory^19,20^, fear memory^21,22^, anxiety^23,24^, and social behavior^25,26^. In addition, electrophysiological experiments revealed that ventral HPC not only innervates excitatory neurons, but also inhibitory interneurons in MFC^27–29^, in such a manner that ventral HPC innervation of specific populations of γ-aminobutyric acid (GABA)-ergic neurons in MFC is important for memory^19,21^, emotional regulation^24,2624,26^, and social behavior^25^.

Although most studies focused on the ventral HPC projection to MFC, the caudal part of dorsal HPC, which includes interconnected, adjacent parts of CA1 and subiculum, generally referred to as distal CA1 and proximal SUB, also has direct projections to MFC^9,10^. Research sustains that synchronized oscillatory activity between dorsal HPC and MFC is important for memory consolidation and learning^30–32^ and that the direct dorsal-caudal HPC-MFC may play a crucial role in transmitting the contextual information processed in dorsal HPC to MFC^33^ and even enhance fear memories through reconsolidation processes^34,35^. Despite such latent functions, the direct dorsal-caudal HPC-MFC circuit has been largely overlooked, and the detailed structural organization of this pathway needs yet to be elucidated.

Here, we examined the organization of the direct dorsal-caudal HPC-MFC circuit by using an anterograde transsynaptic tracing method with adeno-associated viral vectors (AAV) and compared it with that of the ventral HPC-MFC circuit. Our results revealed that dorsal-caudal and ventral HPC target different MFC subregions: ventral HPC targets mainly IL, ventral part of PL, and medial orbital (MO) cortices, whereas dorsal-caudal HPC targets preferentially the most ventral MFC subregions, DP and MO cortices. We also found that about 40% of the neuron population receiving HPC inputs corresponded to GABAergic interneurons, with dcHPC innervating a significant larger proportion of MFC GABAergic interneurons than vHPC. Furthermore, whereas vHPC innervated similar proportions of PV+ and SOM+ interneurons, dcHPC recruited a larger proportion of PV+ interneurons in MFC. Overall, our findings revealed two parallel HPC-MFC circuits, differently organized along the dorsoventral axis, which not only targeted different subregions in MFC, but also engaged distinct proportions of excitatory and inhibitory neurons in MFC.

## RESULTS

### Dorsal-caudal and ventral HPC project differently to MFC subregions in mice

To re-examine the projections from HPC to MFC, anterograde tracers Phaseolus vulgaris-leucoagglutinin (PHA-L) or biotin dextran amine (BDA) were injected along the longitudinal axis of HPC in mice (10 animals, **Figure 1**). According to their dorsoventral position in HPC, the injection sites were clustered in four groups: the dorsal portion of HPC (dHPC, n=3), the dorsal-caudal portion of HPC (dcHPC, n=4), the intermediate to ventral levels of HPC (ivHPC, n=5), and the ventral HPC (vHPC, n=4) (**Figure 1A**). Though in general, projections from HPC innervated caudal portions of MFC more densely, the four groups presented characteristic projection patterns to different subregions of MFC. Injections in dHPC, predominantly located in the proximal CA1 of anterior sections of HPC, did not result in any clear, distinguishable label in MFC (**Figure 1B**, *top-left*). This is in line with previous reports^9,10^. Injections in dcHPC, involving the distal portion of CA1 and the proximal portion of SUB, resulted in dense labeling in all layers of DP and MO, spreading lightly to deep layers of IL, PL, and ACd (**Figure 1B**, *bottom-left*). This particular topographical organization of the projections from dcHPC was not clearly described in previous studies^9,10^. Injections in ivHPC, located more ventrally, mainly involving intermediate parts of CA1, resulted in labeled fibers distributed densely throughout all layers of MO and IL, moderately to the ventral part of PL (PLv) and DP, spreading to deep layers of the dorsal part of PL (PLd) (**Figure 1B**, *top-right*). Finally, injections in vHPC, involved more ventral and posterior sections of field CA1, and in some cases included ventral SUB. The projection pattern observed following these vHPC injections was consistent with previous studies, with fibers spreading densely into all layers of MO, IL, and PLv, and moderately into deep layers of DP and PLd (**Figure 1B**, *bottom-right*)^9,10^.

**Figure 1.**
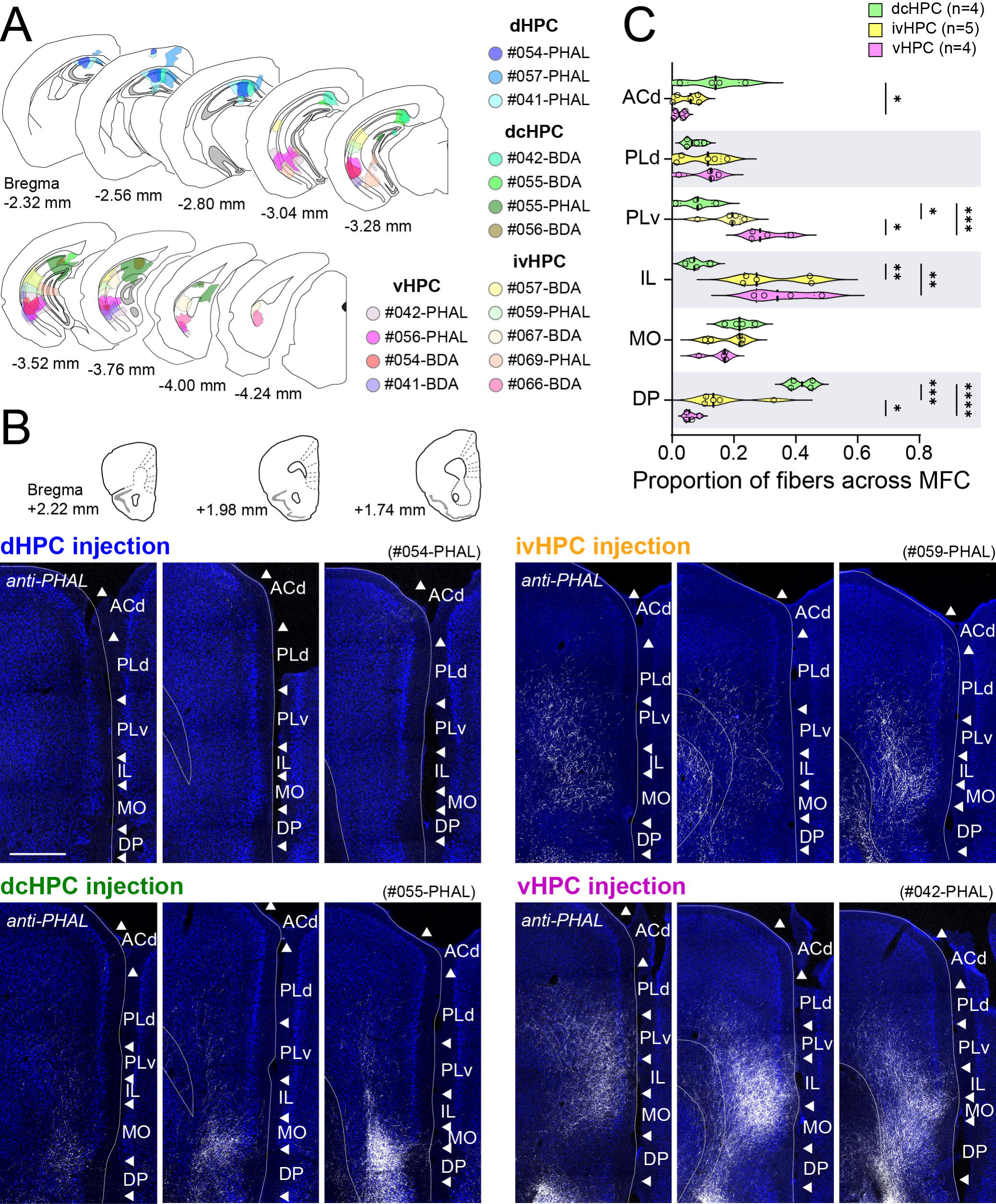
Projections originating along the dorsoventral axis of HPC, visualized by anterograde tracer injections, are differently distributed across MFC subregions in mice. (A) Summary of anterograde tracer (BDA or PHA-L) injection sites along the dorsoventral axis of HPC displayed in coronal sections. HPC injection level was differentiated into dorsal (dHPC), dorsal-caudal (dcHPC), intermediate-ventral (ivHPC), and ventral HPC (vHPC). (B) Representative samples of anterogradely labeled fibers in MFC (at three coronal levels as shown in *top* inset) for injections in dHPC (case #054-PHAL), dcHPC (case #055-PHAL), ivHPC (case #059-PHAL), or vHPC (case #042-PHAL). Scale bar, 500 µm. (C) Quantification of the proportion of labeled fibers across MFC subregions for injections in dcHPC (green, n=4), ivHPC (yellow, n=5), and vHPC (pink, n=4). Each circle represents the proportion of labeled fibers in the MFC subregion among the total observed in MFC for one injection. Values from all subregions sum up to 1 for one individual sample. Data are presented as violin plots. One-way ANOVA, followed by Holm-Šídák’s multiple comparisons test: ****p<0.0001, ***p<0.001; **p<0.01; *p<0.05.

Since the distribution patterns did not differ between the samples with pure CA1 injection and that with CA1+SUB injection, these samples were grouped together in the following analysis. To verify the differences in the organization of these parallel HPC-MFC pathways, we calculated the proportion of labeled fibers detected in each MFC subregion among the total observed in MFC (**Figure 1C**) and analyzed whether the projection patterns from distinct levels of HPC differed significantly in the targeted MFC subregions. In view of the sparse labeling observed in MFC from dHPC injections (**Figure 1B**, *top-left*), this group was excluded from further analysis. The results showed that PLd and MO subregions were targeted similarly by all projecting HPC levels (for PLd, vHPC vs ivHPC, p=0.79; vHPC vs dcHPC, p=0.77; ivHPC vs dcHPC, p=0.77; for MO, vHPC vs ivHPC, p=0.48; vHPC vs dcHPC, p=0.22; ivHPC vs dcHPC, p=0.48; one-way ANOVA followed by Holm-Šídák’s multiple comparison test). Both IL and PLv received stronger innervations from ivHPC and vHPC (for IL, vHPC vs ivHPC, p=0.63; vHPC vs dcHPC, p=0.004; ivHPC vs dcHPC, p=0.004; for PLv, vHPC vs ivHPC, p=0.02; vHPC vs dcHPC, p=0.0008; ivHPC vs dcHPC, p=0.02). In contrast, ACd and DP received denser innervation from dcHPC (for ACd, vHPC vs ivHPC, p=0.49; vHPC vs dcHPC, p=0.046; ivHPC vs dcHPC, p=0.08; for DP, vHPC vs ivHPC, p=0.03; vHPC vs dcHPC, p<0.0001; ivHPC vs dcHPC, p=0.0003). Together, these results revealed that although dorsal-to-ventral HPC levels innervate all MFC subregions, projections from HPC to MFC are organized along the dorsoventral axis. Injections in ivHPC presented a projection pattern quite alike vHPC injections with IL cortex as their main target. Conversely, dcHPC injections presented a significant difference in main targeted MFC subregions, heavily innervating the more ventrally located DP cortex and less strongly ACd. Thus, in the subsequent sections we will focus on comparing the parallel pathways from dcHPC and vHPC to MFC.

### dcHPC and vHPC project differently to MFC subregions in rats

One reason that the difference between the dcHPC-MFC and vHPC-MFC projection patterns has not been clearly addressed in previous studies could be the species differences. Although previous studies used rats^9,10^, we used mice in the above experiment. To confirm the consistency of the HPC-MFC circuits across rodent species, we conducted the same tracing experiments in rats (**Figure S1**). Similar to the results in mice (**Figure S1A**), after tracer injections in dHPC, labeling in MFC was weak to absent (**Figure S1B**). Injections in dcHPC resulted in very dense fiber-labeling in DP, the most ventral MFC subregion (**Figure S1C**). Conversely, injections in vHPC resulted in dense labeled fibers in IL and PLv subregions (**Figure S1D**). These data thus indicate that the projection patterns of HPC-MFC pathways is similar between mice and rats.

In contrast to many cortico-cortical connections that are reciprocal, a direct MFC-HPC projection is apparently absent. Several studies have shown that MFC is indirectly connected to HPC via either midline thalamic nuclei or the entorhinal cortex (EC)^36–38^. Interestingly, a retrograde tracing study using pseudorabies virus showed that these indirect pathways show a topographical organization along the dorsoventral axis of HPC^39^. In the latter study it was reported that two parallel pathways connect the frontal cortex with HPC, a dorsal pathway from AC to dHPC through dorsolateral EC or the anterior thalamus, and a ventral pathway from ventral MFC to vHPC, through caudomedial EC or the midline thalamus. To compare the projection patterns of the direct HPC-MFC pathway with that of the indirect MFC-HPC pathway, we examined the organization of multisynaptic MFC inputs to dcHPC and vHPC. We conducted a retrograde transsynaptic tracing experiment with rabies virus (RV) in rats. We first examined the brain regions which have direct projections to HPC (7 animals) by using either a glycoprotein-deleted RV vector (ΔG -RV), which can be used as a non-transsynaptic retrograde tracer^40^, or a propagation-competent RV vector, labeling 1^st^ order projection neurons with two days of survival time^41^ (**Figure 2A**). Irrespective of the RV vector used, many retrogradely labeled neurons were observed in the thalamic nucleus reuniens (Re) and layer II and III of EC with topographical labeling patterns depending on the position of the injection site along the HPC longitudinal axis (**Figure 2C1-2C2**). The RV vector injected into dcHPC (n=4) resulted in labeled neurons positioned medially in Re and dorsally in lateral EC (LEC), whereas the neurons labeled by the RV vector injected into vHPC (n=6) were preferentially found laterally in Re and ventrally in LEC. In all of the cases, only very few labeled neurons were observed in MFC which corroborates previous studies that MFC does not, or only very sparsely, projects directly to HPC^42–44^ (**Figure 2C3, 2E, 2G**).

**Figure 2.**
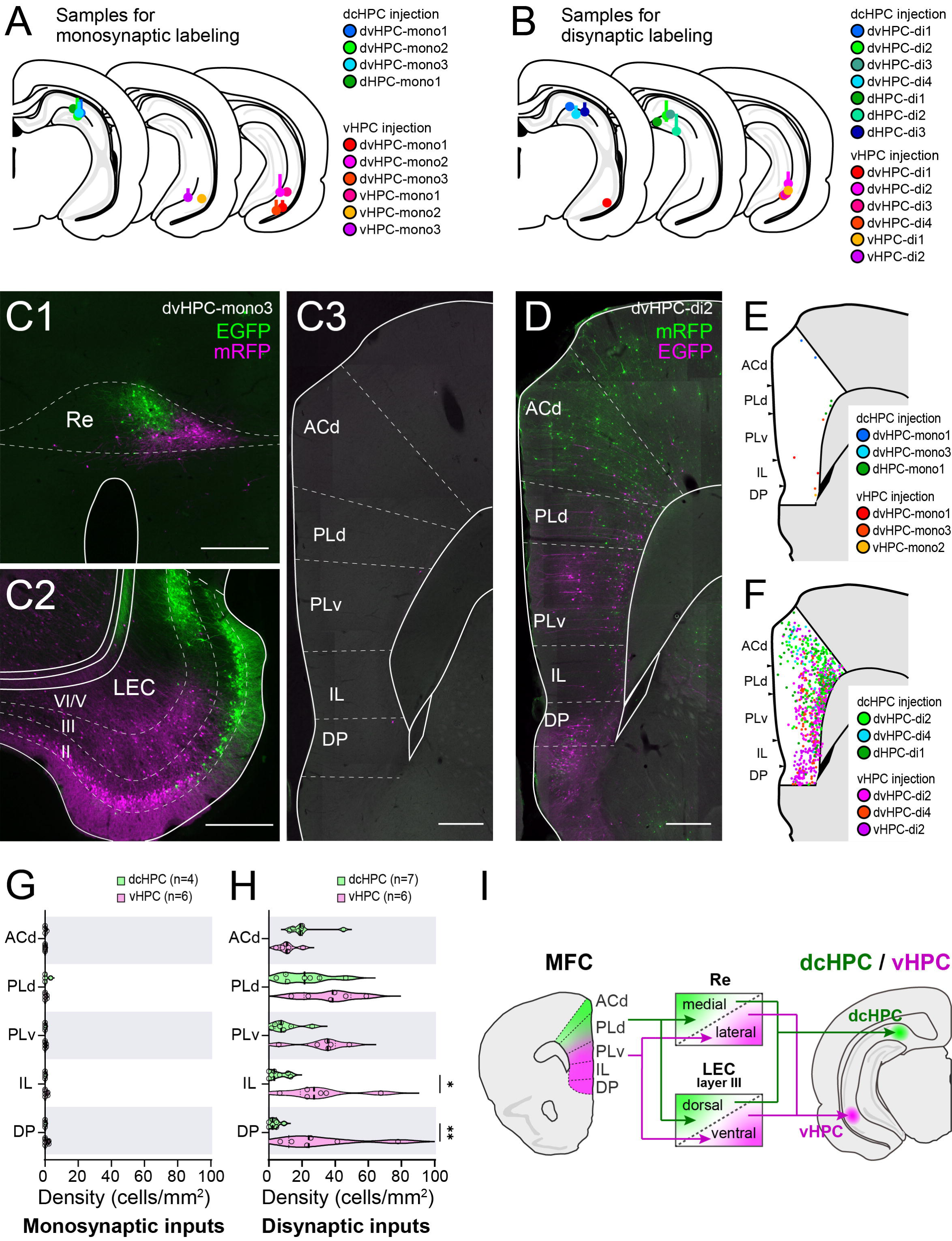
Organization of the disynaptic inputs from MFC to HPC along the dorsoventral axis. (A-B) Summary of injection sites of RV vectors in dcHPC and vHPC. Each injection is illustrated with a different color. Either a glycoprotein-deleted RV vector or a propagation-competent RV vector with short survival time was used to examine monosynaptic inputs to dcHPC (n=4) and vHPC (n=6) (A). A propagation-competent RV vector with longer survival time was used to examine disynaptic inputs to dcHPC (n=7) and vHPC (n=6) (B). (C) Fluorescence micrograph of retrograde labeling in Re (C1), LEC (C2), and MFC (C3) of a monosynaptically labeled sample (case dvHPC-mono3). Neurons infected by the RV vector injected to dcHPC are shown in green, while neurons infected by the vHPC-injected vector are shown in magenta. Scale bars, 500 µm. (D) Fluorescence micrograph of retrograde labeling in MFC of a disynaptically-labeled sample (case dvHPC-multi2). Scale bar, 500 µm. (E-F) Distribution of labeled MFC neurons from three representative samples are overlaid for monosynaptic (E) and disynaptic samples (F) for injections in dcHPC (cool colors) and vHPC (warm colors). (G-H) Density of labeled neurons across MFC subregions are shown for monosyanptic-(G) and disynaptic-labeling samples (H) after dcHPC (green) and vHPC (pink) injections. Each circle represents the individual density of labeled cells in the subregion for one injection sample. Data are presented as violin plots. Mann-Whitney test corrected for multiple comparisons using the Holm-Šídák method: **p<0.01, *p<0.05. (I) Summary illustration of parallel disynaptic HPC pathways from dorsal and ventral MFC to dcHPC and vHPC, via Re or LEC layer III neurons.

To examine the disynaptic inputs to HPC (9 animals), we next used the propagation-competent RV vector with four to five days of survival time, which are optimal survival times for the viral vector to transsynaptically label the 2^nd^-order projection neurons (**Figure 2B**). In these samples, the number of labeled neurons dramatically increased in MFC, and the two populations of labeled cells exhibited a topographical distribution (**Figure 2D, 2F, 2H**). The neurons labeled by the ventral injection (n=6) were prominent in the ventral part of MFC, including DP, IL, and PLv. In contrast, neurons labeled by the dorsal-caudal injection (n=7) were mainly distributed in the dorsal part of MFC, particularly in ACd. The density of labeled neurons in PLd was similar for dorsal and ventral injections. Interestingly, we did not see such massive MFC labeling when the RV was injected specifically in DG^40^, despite a large number of 1^st^ order labeled cells that were observed in layer II of LEC. Thus, the combined experimental data of this series of tracing experiments indicate that MFC neurons disynaptically project to CA1/SUB either via Re or layer III of EC in a topographical manner along the dorsoventral axis (**Figure 2I**).

Taken together, these rat anatomical data indicate that there is a crosstalk between the indirect MFC-HPC circuit and the direct HPC-MFC circuit. The information from MFC to ventral CA1/SUB, originated in PLd, PLv, IL, and DP, will be sent back mainly to PLv and IL. In contrast, information from MFC to dorsal CA1/SUB, originated mainly in ACd and PLd, will be sent back to DP, the most ventral MFC subregion.

### dcHPC and vHPC elicit excitatory and inhibitory responses in projection neurons in MFC

Projections from the different MFC subregions to amygdala (AMG) are thought to be important in the modulation of anxiety-and fear-related behaviors^45–48^. Inputs from HPC seem to be able to modulate the activity of these AMG-projecting neurons in MFC and influence behavior^21^. Interestingly, our anatomical data revealed that dcHPC and vHPC present significantly distinct projection patterns to MFC, suggesting that their innervation may differentially influence the projection neurons in the MFC subregions. Thus, to confirm the distinct targeting of dcHPC or vHPC inputs to the MFC subregions, we next applied optogenetics and whole-cell recordings of AMG-projecting neurons in MFC in mice (23 animals). First, channelrhodopsin-2 (ChR2) was expressed in either dcHPC or vHPC by AAV transduction, and in parallel, either the retrograde AAVrg-hSyn-mCherry or red Retrobeads were injected in AMG. We then obtained whole-cell voltage-clamp recordings from AMG-projecting neurons in acute coronal slices of MFC, while optogenetically stimulating the terminal axons from dcHPC or vHPC (**Figure 3A**, *left*). Recorded neurons were labeled with biocytin for later verification of their location within the MFC subregions. We assessed the relative excitatory and inhibitory drive by recording light-evoked synaptic input at holding potential of -70 mV for excitatory postsynaptic currents (EPSCs), and at +10 mV for inhibitory postsynaptic currents (IPSCs). Since HPC innervations are known to be of glutamatergic or excitatory nature in MFC^29,49,50^, we assumed that the IPSC responses would be mediated by disynaptic inhibition from local interneurons (**Figure 3A**, *right*).

**Figure 3.**
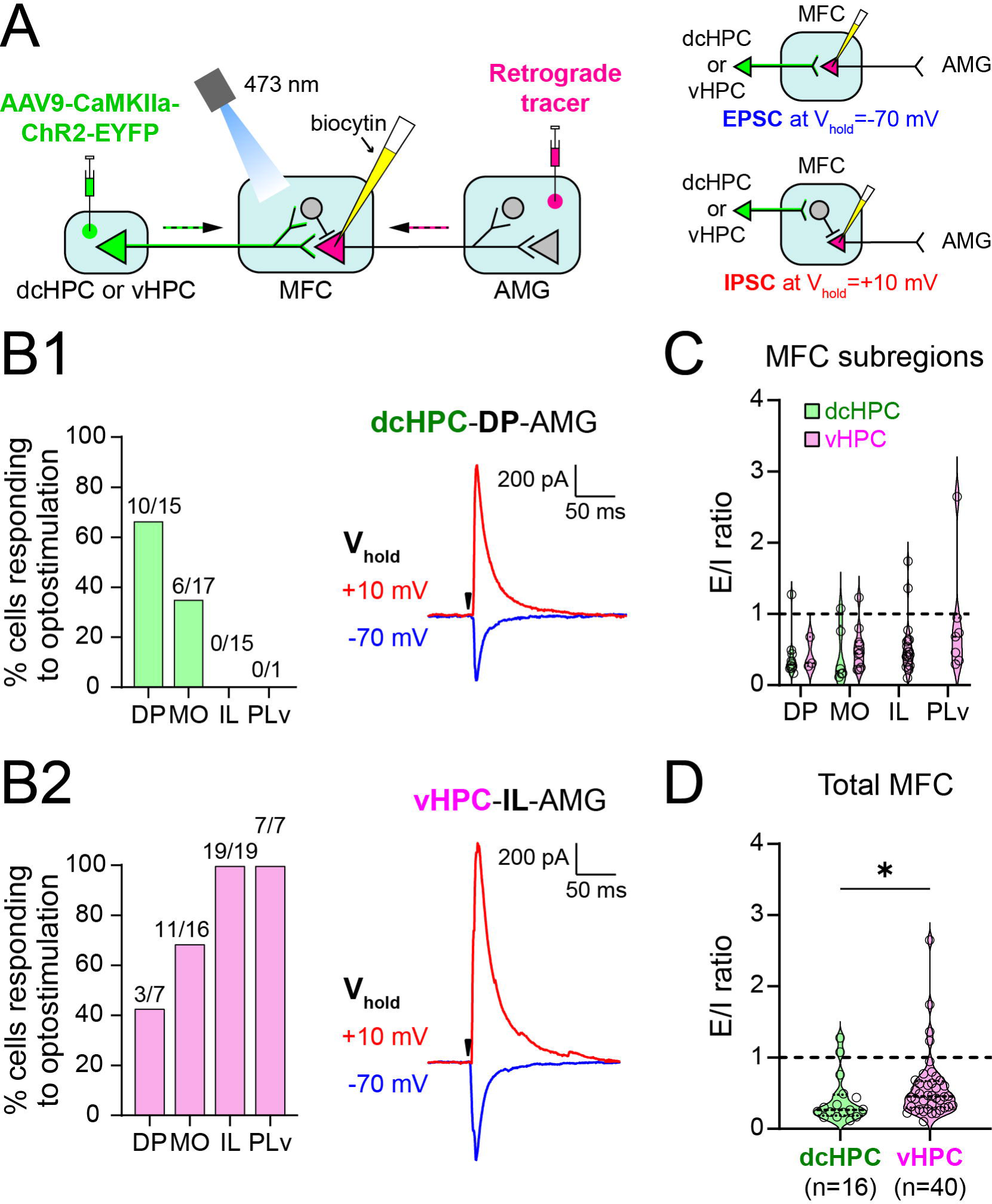
dcHPC and vHPC connect neurons in different subregions in MFC. (A) *Left*, general scheme of the electrophysiological experiment: AAV9-CaMKIIa-ChR2-EYFP was injected in either dcHPC or vHPC to express ChR2 in projection neurons. In parallel, the retrograde AAVrg-hSyn-mCherry, or in some cases red Retrobeads, was injected in AMG to label AMG-projecting neurons in MFC. Whole-cell voltage-clamp recordings were carried out in acute MFC coronal slices from AMG-projecting neurons (rose-colored), while optically stimulating the dcHPC or vHPC axon terminals in MFC. Recorded neurons were labeled with biocytin for posterior visualization. *Right*, strategy used to record MFC excitatory input at holding potential of -70 mV (EPSC, *top*), or inhibitory input (IPSC, *bottom*) mediated via local interneurons at holding potential of +10 mV. In the schemes, principal neurons and interneurons are represented by triangles and circles, respectively. (B1) *Left*, proportion of responding cells to optical stimulation of dcHPC axon terminals across the MFC subregions. *Right*, responses from an AMG-projecting cell in DP to light-evoked dcHPC inputs at holding potential of -70 mV (EPSC, blue) and +10 mV (IPSC, red). Black arrow indicates light pulse. (B2) *Left*, proportion of responding cells to optical stimulation of vHPC axon terminals across the MFC subregions. *Right*, responses from an AMG-projecting cell in IL to light-evoked vHPC inputs at holding potential of -70 mV (EPSC, blue) and +10 mV (IPSC, red). Black arrow indicates light pulse. (C) Summary of the ratio of light-evoked responses recorded at -70 mV and +10 mV presented as an excitatory to inhibitory (E/I) ratio. Higher values mean input is biased to excitation, whereas low values towards inhibition. Cells receiving inputs from dcHPC and vHPC are presented in green and pink, respectively. Data are shown as violin plots, where each circle represents one cell. No significant differences were found across the MFC subregions (p=0.1, Kruskal-Wallis Test). (D) Summary of the light-evoked E/I ratios from all recorded cells in MFC for dcHPC (green, n=16 cells) or vHPC inputs (pink, n=40 cells). Data are presented as violin plots, where each circle corresponds to one cell. Two-tailed Mann-Whitney Test: *p<0.05.

We found that the connectivity between dcHPC and vHPC with the MFC subregions reflected our anatomical observations. Optical stimulation of dcHPC axon fibers evoked responses in AMG-projecting neurons mainly located in DP (66.7%, 10 out of 15 cells) and MO (35.3%, 6 out of 17 cells), whereas no recorded neuron responded in IL (0 out of 15 cells) and PLv (0 out of 1 cell) (**Figure 3B1**, *left*). In contrast, the light stimulation of vHPC axon fibers evoked responses in all recorded AMG-projecting neurons in IL (19 out of 19 cells) and PLv (7 out of 7 cells), and to a lesser extent in MO (68.8%, 11 out of 16 cells) and DP (42.9%, 3 out of 7 cells, **Figure 3B**2, *left*). The optical stimulation of either dcHPC or vHPC terminals evoked excitatory and inhibitory responses in the AMG-projecting neurons, recorded at -70 mV (EPSC) and at +10 mV (IPSC) respectively (**Figure 3B1**, *right*, **3B2**, *right*). The mean latencies of the EPSC and IPSC responses were significantly different, with IPSC lagging EPSC when optically stimulating the vHPC fibers (EPSC, 5.1±0.3 ms; IPSC, 7.3±0.3 ms; mean±SEM, p<0.0001, two-tailed t-test) and the dcHPC fibers in MFC (EPSC, 5.6±0.4 ms; IPSC, 8.7±0.7 ms; mean±SEM, p=0.0003, two-tailed t-test). Furthermore, in both vHPC and dcHPC cases, IPSCs, but not EPSCs, were abolished when isolating the monosynaptic inputs by perfusing the recording solution with a mix of TTX (1 µM), 4-AP (100 µM), and elevated Ca^2+^ (4 mM). IPSCs were also abolished by applying gabazine (10 µM) to the extracellular solution, which blocks the GABA-_A_ receptors (*data not shown*). These observations indicated that, as proposed above (**Figure 3A**, *right*) and in line with the postulated glutamatergic nature of the HPC inputs to MFC^29,49,50^, the evoked IPSC responses may be indeed a consequence of the HPC innervation of the inhibitory local circuit in MFC. By comparing the ratio of excitatory to inhibitory (E/I) responses from the responsive cells, we observed that inputs from both dcHPC and vHPC seem to drive substantial disynaptic inhibition in AMG-projecting neurons, with no significant differences within the MFC subregions (**Figure 3C**, p=0.46, Kruskal-Wallis Test). Interestingly, when the E/I ratios from the total of responsive cells in MFC were analyzed as a group for each HPC case, vHPC seemed to have an overall stronger excitatory drive over the AMG-projecting neurons in MFC than dcHPC (**Figure 3D**; vHPC=0.45, n=40 cells; dcHPC=0.27, n=16 cells; p=0.03, two-tailed Mann-Whitney Test).

These results confirm that dcHPC and vHPC axons contact neurons in different subregions of MFC in accordance with the projection patterns described in our tracing experiments, i.e. dcHPC mainly engages neurons located in DP and MO, whereas vHPC innervates neurons in all MFC subregions, but with a substantial preference for neurons located in IL and PLv. Furthermore, these two parallel pathways presented different overall ratios of excitatory and inhibitory influence on MFC, suggesting that dcHPC and vHPC may be innervating different proportions of excitatory and inhibitory neurons in MFC.

### Distribution of postsynaptic neurons in MFC subregions targeted by dcHPC and vHPC

To identify the postsynaptic neurons receiving HPC inputs in MFC, we next used an anterograde transsynaptic tagging strategy with adeno-associated virus serotype 1 (AAV1) in mice (11 animals) (**Figure 4A**). AAV1 can spread anterogradely to the postsynaptic neurons when injected at a high titer^51^. Although AAV1 can also transport retrogradely at high titer, this is not of concern in this experiment since these two regions are, mainly unidirectionally connected through the direct projections from HPC to MFC^52^. Here, we injected a Cre-expressing AAV1 (AAV1-hSyn-Cre) in dcHPC (n=6) or vHPC (n=5) (**Figure 4B**) and identified the postsynaptic neurons in MFC, either by immunostaining against the Cre-recombinase (**Figure 4**) or by the presence of fluorescent protein expressed via a Cre-dependent-AAV injected in MFC (**Figure 5, 6**). In line with the observations of our anterograde tracing experiments (**Figure 1**), dcHPC innervated mainly neurons in DP and MO, whereas vHPC innervated mainly neurons in IL, PLv, and MO (**Figure 4C**). To describe the distribution of the neurons receiving HPC inputs across the MFC subregions, we quantified a subregional proportion as the number of Cre-labeled cells in a subregion among the total number of Cre-labeled cells in MFC. The distribution of HPC postsynaptic cells across the MFC subregions (**Figure 4D**) presented comparable patterns to the ones observed in our previous experiment using conventional anterograde tracing (**Figure 1C**). Compared to dcHPC, projections from vHPC innervated a larger proportion of neurons in IL and PLv (dcHPC vs vHPC; IL, p=0.03; PLv, p=0.03; Mann-Whitney test). In contrast, dcHPC presented a larger proportion of inputs into DP than vHPC (dcHPC vs vHPC; p=0.03, Mann-Whitney test). The proportion of cells receiving inputs from dcHPC or vHPC were similar in ACd, PLd, and MO (dcHPC vs vHPC; ACd, p=0.2; PLd, p=0.4; MO, p>0.99; Mann-Whitney test). Although we did not look into the distribution of Cre-labeled cells along the anterior-posterior axis of MFC, it seems that whereas vHPC targets IL and PLv densely in middle sections of MFC, dcHPC projections seem denser in DP in middle-to-posterior sections of MFC **(Figure S2)**.

**Figure 4.**
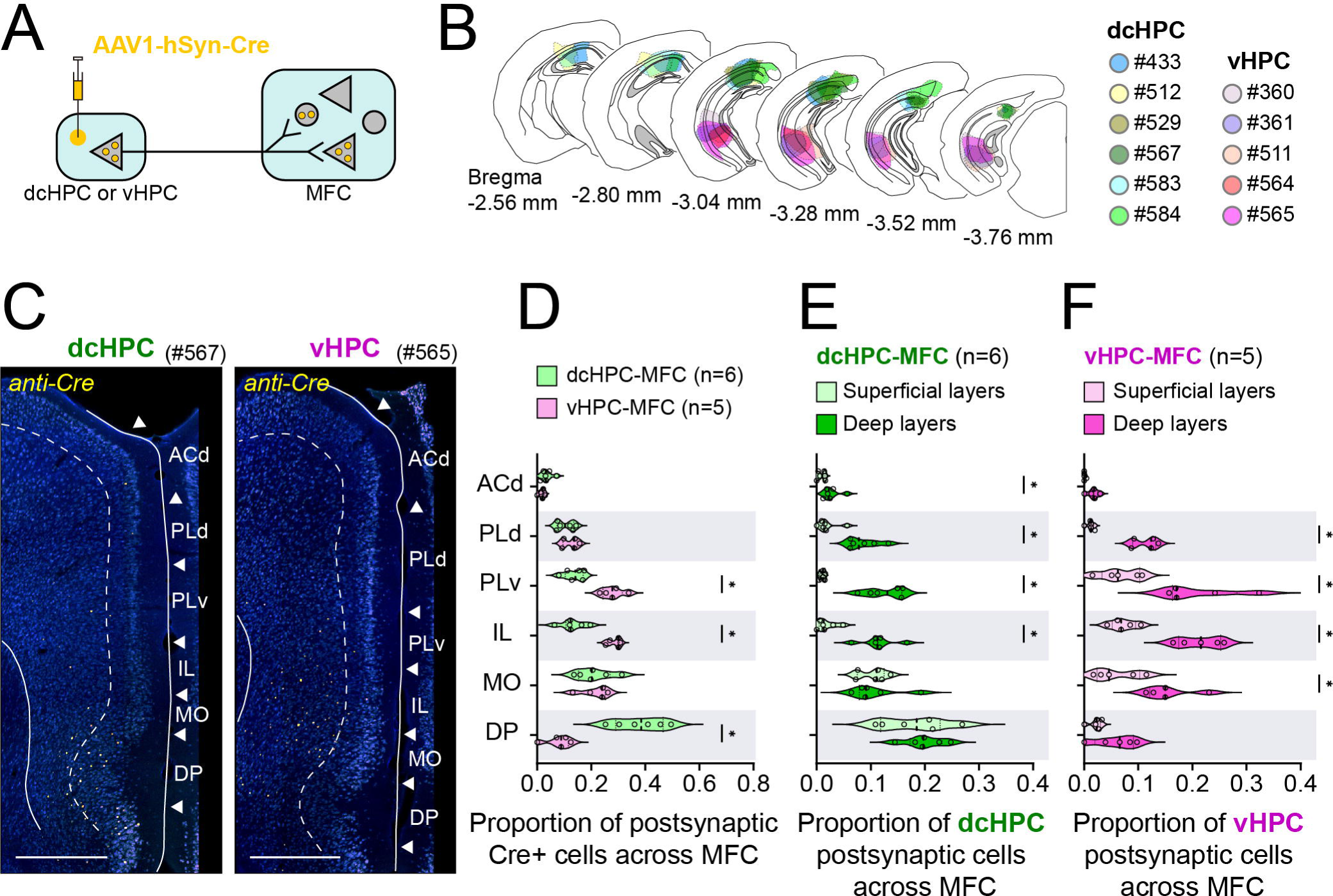
Anterograde transsynaptic spread of AAV1-Cre injections from dcHPC and vHPC target superficial and deep layers of MFC. (A) AAV1-hSyn-Cre was injected in either dcHPC or vHPC resulting in expression in HPC projection neurons (not shown). The transsynaptic spread to postsynaptic neurons was detected by immunostaining against Cre. Note that postsynaptic neurons likely include principal neurons and interneurons in MFC which in the scheme, are represented by triangles and circles, respectively. The present experiments do not include a further differentiation between interneurons and principal neurons. (B) Summary of AAV1-Cre injection sites along HPC displayed in coronal sections. Injection sites in dcHPC and vHPC are shown in cool and warm colors, respectively. (C) Representative samples of MFC coronal sections showing the distribution of postsynaptic Cre-expressing neurons in MFC for dcHPC (*left*, case #567) and vHPC (*right*, case #565) injections. Scale bars, 500 µm. (D) Proportion of the number of Cre-labeled cells in a MFC subregion among the total Cre-labeled neurons across MFC sections for dcHPC (green, n=5) and vHPC (magenta, n=5) injections. Data are presented as violin plots, with each circle corresponding to one sample. Values from all subregions sum up to 1 for one individual sample. Mann-Whitney test corrected for multiple comparisons using the Holm-Šídák method: *p<0.05. (E, F) Laminar distribution of postsynaptic Cre-labeled cells in superficial and deep layers of MFC subregions for dcHPC (E) and vHPC (F) injections. Data are presented as violin plots, with each circle corresponding to one sample. Values from superficial and deep layers of all subregions sum up to 1 for one individual sample. Mann-Whitney test corrected for multiple comparisons using the Holm-Šídák method: *p<0.05.

**Figure 5.**
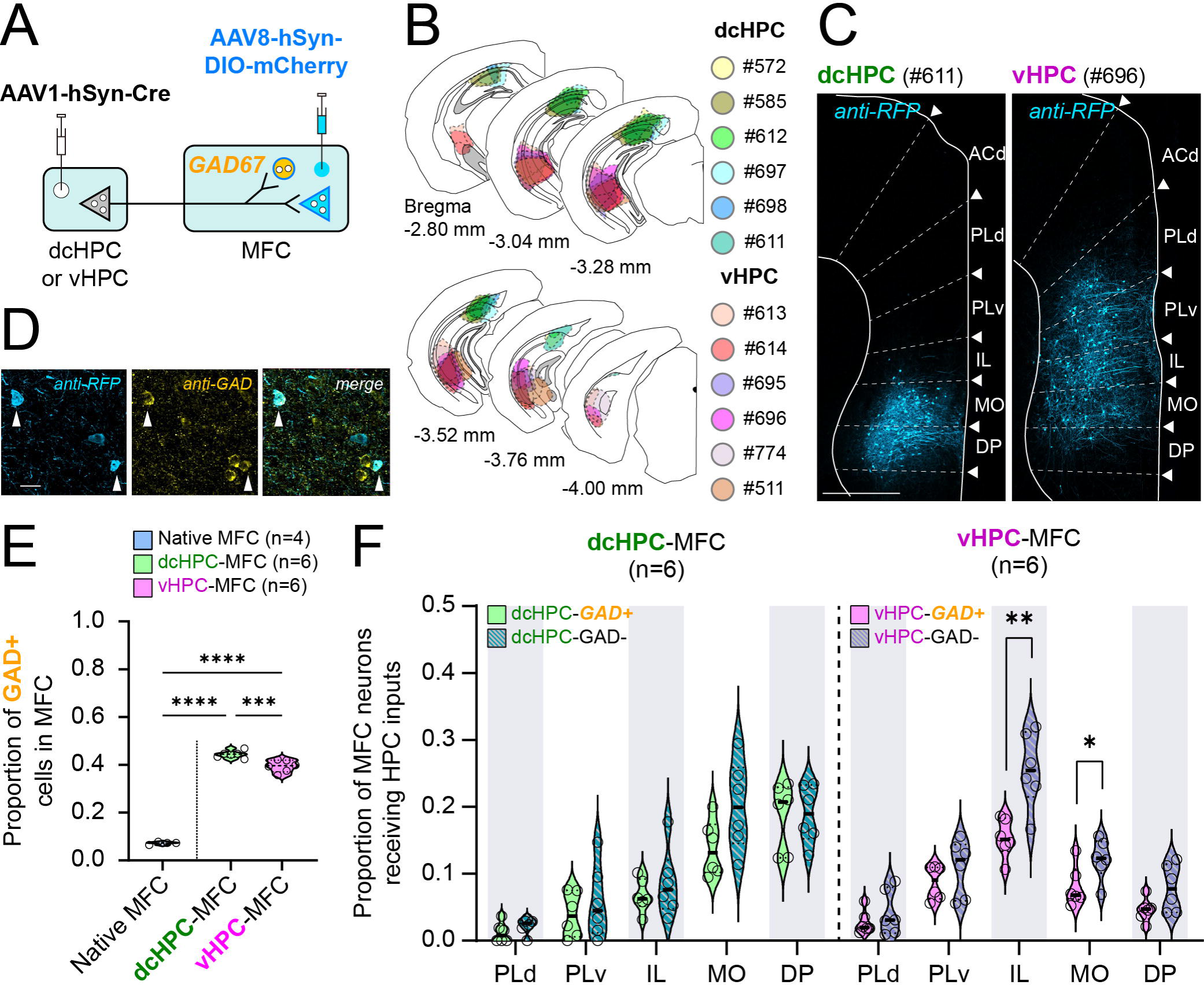
dcHPC and vHPC inputs to MFC innervate GABAergic and non-GABAergic neurons. (A) AAV1-hSyn-Cre was injected in either dcHPC or vHPC and was expressed in HPC projection neurons. The transsynaptic spread led to Cre-expression in postsynaptic neurons, including principal neurons and interneurons in MFC. In parallel, AAV8-hSyn-DIO-mCherry was injected in MFC leading to Cre-dependent mCherry-expression in postsynaptic neurons in MFC. Postsynaptic interneurons were subsequently differentiated by immunostaining against GAD67, as illustrated in D-F. In the scheme, principal neurons and interneurons are represented by triangles and circles, respectively. (B) Summary of AAV1-hSyn-Cre injection sites along the HPC displayed in coronal sections. Injection sites in the dcHPC and vHPC are shown in cool and warm colors, respectively. (C) Representative samples with AAV1-hSyn-Cre injection in either dcHPC (*left*, case #611) or vHPC (*right*, case #696). Images show the postsynaptic Cre-dependent-mCherry-expression in a coronal section of MFC. Scale bar, 500 µm. (D) Representative sample showing the mCherry-expression of postsynaptic neurons and colocalization with the GAD67 antibody labeling as used to identify the GABAergic neurons among the HPC-MFC postsynaptic cells. Scale bar, 20 µm. (E) Proportion of cells co-expressing mCherry and GAD67 among the total mCherry+ neurons receiving dcHPC (green, n=6) or vHPC (pink, n=6) inputs in MFC. The proportions were compared to the native presence of GAD67 in MFC (light-blue, n=4). One-way ANOVA, followed by *post hoc* Tukey’s multiple comparison test: ***p<0.001, ****p<0.0001. (F) Subregional distribution of GABAergic (GAD+) and non-GABAergic (GAD-) neurons in MFC receiving inputs from dcHPC (*left*, n=6, green tones) and vHPC (*right*, n=6, pink tones) Cre-injections. The proportions of inputs to GAD+ and GAD-cells across MFC subregions sum up to 1 for each injection sample. Data is presented as violin plots, where each circle corresponds to one sample. Two-tailed t-tests: **p<0.01, *p<0.05.

We also noticed that the distribution of Cre-expressing cells differed across layers. In general, deep layers (layers V/VI) of MFC presented larger proportions of Cre-labeled cells than superficial layers (layers I/II/III) of MFC. The proportion of neurons receiving inputs from dcHPC (**Figure 4E**) was larger in deep layers than in superficial layers in ACd, PLd, PLv, and IL (deep vs sup; ACd, p=0.04; PLd, p=0.01; PLv, p=0.01; IL, p=0.01; Mann-Whitney test). In MO and DP, however, Cre-labeled cells were similarly distributed across superficial and deep layers (deep vs sup; MO, p=0.9; DP, p=0.9; Mann-Whitney test). Regarding the proportion of labeled neurons receiving inputs from vHPC (**Figure 4F**), we observed that this was larger in deep layers than superficial layers in MO, IL, PLv, and PLd (deep vs sup; MO, p=0.047; IL, p=0.047; PLv, p=0.047; PLd, p=0.047; Mann-Whitney test), whereas the proportions of labeled neurons for deep and superficial layers in ACd and DP were similar (deep vs sup; ACd, p=0.09; DP, p=0.1; Mann-Whitney test). These observations confirm that dcHPC and vHPC project differently to MFC; dcHPC mainly innervates neurons in superficial and deep layers of DP, whereas vHPC sends mainly inputs to deep layers of IL and PLv.

### dcHPC and vHPC innervate significant proportions of GABAergic neurons in MFC

Previous studies have reported that projections from vHPC to MFC play a crucial role in regulating memory and emotional-related behavior through the recruitment of GABAergic neurons in MFC^21,24,25,53^. Interestingly, the results from our whole-cell recordings suggested that the overall E/I ratio in MFC tends to be more inhibitory upon optogenetic stimulation of axon fibers originating in dcHPC than in vHPC (**Figure 3D**). To verify whether dcHPC and vHPC innervate different proportions of GABAergic and non-GABAergic neurons in MFC, we used the anterograde transsynaptic tagging method to label the neurons receiving HPC inputs in MFC (12 animals): AAV1-hSyn-Cre was injected into either dcHPC (n=6) or vHPC (n=6), and in parallel, a Cre-dependent AAV expressing mCherry (AAV8-hSyn-DIO-mCherry) was injected into MFC (**Figure 5A, B**). Consistent with the labeling patterns observed in the above sections, for dcHPC injections, mCherry+ cells were localized mainly in DP and MO (**Figure 5C**, left) whereas for vHPC injections, mCherry+ cells were spread along MO, IL and PLv subregions (**Figure 5C**, right). A visual inspection of the mCherry+ neuron axon terminals throughout the brain (**Figure S3**), showed that MFC neurons receiving inputs from vHPC and dcHPC (**Figure S3A-B**) target the claustrum, insular cortex, septum, nucleus accumbens, mediodorsal thalamus, basal amygdala, hypothalamus, ventral tegmental area, and dorsal raphe nucleus (**Figure S3C-M**). These observed projections are in line with previous anatomical studies on the efferent projections from MFC^54,55^. Interestingly, MFC neurons receiving dcHPC inputs, target distinctively the mammillary body (**Figure S3L**), ventral tegmental area (**Figure S3K**) and submedial nucleus of the thalamus (**Figure S3I**).

Next, by immunostaining against glutamic acid decarboxylase (GAD67), the mCherry+ cells were differentiated as GABAergic (GAD+) or non-GABAergic (GAD-) neurons (**Figure 5A, 5D**). The proportions of GAD+ neurons among the total of mCherry+ cells in MFC receiving inputs from dcHPC or vHPC were calculated and then compared with the native proportion of GAD+ cells in MFC, corresponding to the overall proportion of GABAergic neurons present in MFC (**Figure 5E**). Both dcHPC and vHPC projections innervated a significant large proportion of GAD+ cells considering the native presence of GAD+ cells in MFC (native MFC, 0.072±0.006; dcHPC-MFC, 0.44±0.01; vHPC-MFC, 0.39±0.02; mean±SD, F_(2,13)_=543.3, p<0.0001, one-way ANOVA; dcHPC-MFC vs native MFC, p<0.0001; vHPC-MFC vs native MFC, p<0.0001; one-way ANOVA followed by *post hoc* Tukey’s multiple comparison test). Interestingly, a larger proportion of dcHPC postsynaptic neurons in MFC were GAD+ than among vHPC postsynaptic neurons (dcHPC-MFC vs vHPC-MFC, p=0.0005; one-way ANOVA followed by *post hoc* Tukey’s multiple comparison test).

To further characterize the dcHPC-MFC and vHPC-MFC circuits, we calculated the proportions of GAD+ and GAD-neurons among the total mCherry+ cells receiving HPC inputs for each MFC subregion (**Figure 5F**). Because mCherry-expression was absent in ACd, this subregion was excluded from this analysis. Projections from dcHPC innervated similar proportions of GAD+ and GAD-cells in the MFC subregions (GAD+ vs GAD-; PLd, p=0.21; PLv, p=0.48; IL, p=0.44; MO, p=0.07; DP, p=0.95; two-tailed t-test). In contrast, vHPC innervated similar proportions of GAD+ and GAD-cells in MFC subregions with the exception of IL and MO (GAD+ vs GAD-; PLd, p=0.38; PLv, p=0.29; DP, p=0.07; two-tailed t-test). In IL and MO, a larger proportion of GAD-than GAD+ neurons received vHPC inputs (IL, p=0.003; MO, p=0.04, two-tailed t-test), which suggested that vHPC favors excitatory innervation in IL and MO. These results show that both dcHPC and vHPC innervate significant proportions of GABAergic cells in the MFC subregions, and furthermore, they suggest that vHPC may exert a stronger excitatory effect in IL and to a less extent in MO.

### dcHPC and vHPC recruit parvalbumin- and somatostatin-expressing cells in MFC

Our anatomical observations led to the conclusion that GABAergic neurons are among the monosynaptically innervated neurons in MFC. Previous reports are in line with this and further suggest that vHPC projections may regulate the network dynamics in MFC through the recruitment of different types of inhibitory interneurons found in MFC, including parvalbumin-(PV+), somatostatin- (SOM+), vasoactive intestinal peptide-(VIP+) and cholecystokinin-(CCK+) expressing neurons^53,56,57^.

Two of the main populations of GABAergic neurons in MFC are PV+ and SOM+ interneurons, which have characteristic physiological and morphological properties, and display particular lamination patterns in MFC^53^. Fast-spiking PV+ interneurons target cell bodies and proximal dendrites of pyramidal neurons leading to a high level of feedforward and feedback inhibition^58^. In contrast, SOM+ interneurons form inhibitory synapses on distal dendritic branches of principal neuron enhancing the selectivity of excitatory inputs^59,60^, and leading to long-lasting and delayed inhibition^61^. SOM+ neurons also innervate nearby PV+ cells, thus decreasing PV-mediated inhibition^62^. Interestingly, immunostained sections of MFC against PV and SOM showed that PV+ neurons are densely localized in the most dorsal and most ventral parts of MFC (**Figure 6A**, *left*) whereas SOM+ interneurons seemed to be uniformly distributed across the MFC subregions (**Figure 6A**, *right*). We thus, hypothesized that dcHPC and vHPC innervate different interneuron types in MFC. To examine this, we used an anterograde transsynaptic tagging strategy (**Figure 6B**) in which the AAV1-hSyn-Cre was injected in either dcHPC (n=6) or vHPC (n=4) (**Figure 6C**) and in parallel the AAV9-hDlx-Flex-GFP was injected in MFC (10 animals). The latter viral vector expresses a Cre-dependent-GFP reporter under the control of the hDlx promoter to specifically tag GABAergic neurons as previously reported^63^, which we confirmed by immunostaining against GAD67 (**Figure 6D**). Subsequently, the GFP-expressing neurons were characterized by immunostaining against the markers PV and SOM (**Figure 6E**). We then calculated the proportions of PV+ and SOM+ interneurons among the total GFP-labeled interneurons in MFC. Our results showed that dcHPC and vHPC innervated different proportions of PV+, SOM+ and other kinds of interneurons in MFC (**Figure 6F-G**). Among the interneurons receiving dcHPC inputs, the PV+ proportion was larger compared to the SOM+ and other non-PV/SOM interneurons (PV+ vs SOM+, p=0.024; PV+ vs Others, p=0.021; SOM+ vs Others, p=0.997; one-way ANOVA followed by Tukey’s multiple comparisons test). In contrast, among the interneurons receiving vHPC inputs, the proportions of PV+ and SOM+ were not significantly different (PV+ vs SOM+, p=0.079; PV+ vs Others, p=0.484; SOM+ vs Others, p=0.012; one-way ANOVA followed by Tukey’s multiple comparisons test). Furthermore, we analyzed the subregional distribution of PV+ and SOM+ interneurons receiving HPC inputs in MFC (**Figure 6H-I**). Among the interneurons receiving dcHPC inputs, PV+ and SOM+ cells were mainly located in DP and MO (**Figure 6H**). In contrast, PV+ and SOM+ cells receiving vHPC inputs were mainly located in PLv and IL (**Figure 6I**). Note, however, that we did not find any significant differences in the proportions of postsynaptic PV+ and SOM+ cells within any MFC subregion for either dcHPC **(Figure 6H)** or vHPC **(Figure 6I)** cases.

**Figure 6.**
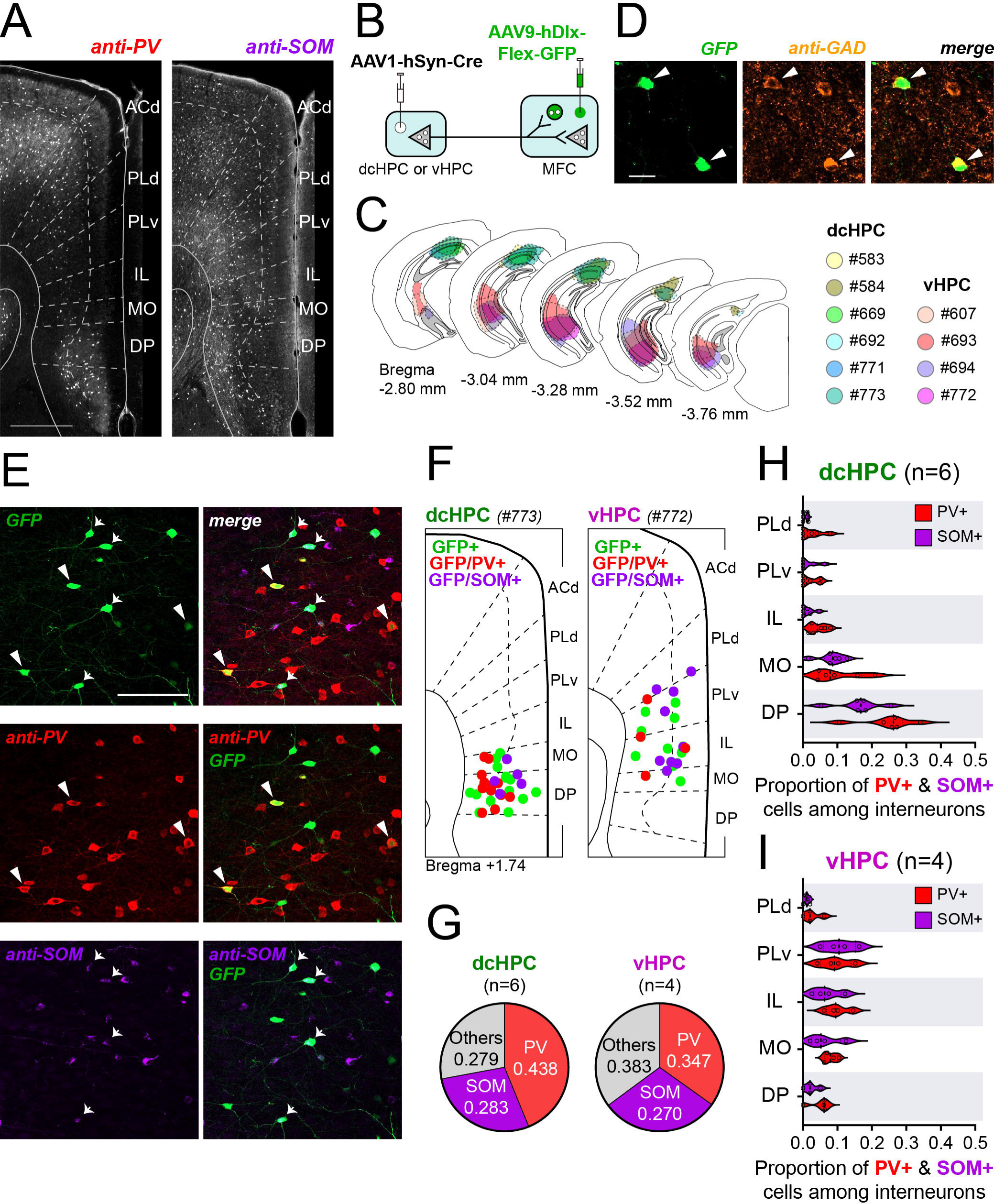
dcHPC and vHPC innervate both PV and SOM interneurons in MFC. (A) Representative images showing the native distribution of the indicated marker expression (*left*, PV; *right*, SOM) in the subregions of MFC. Scale bar, 500 µm. (B) AAV1-hSyn-Cre was injected in either dcHPC or vHPC and expressed in HPC projection neurons. The transsynaptic spread led to Cre-expression in postsynaptic neurons, including principal neurons and interneurons in MFC. In parallel, AAV9-hDlx-Flex-GFP was injected in MFC, leading to Cre-dependent GFP-expression in postsynaptic interneurons in MFC. In the scheme, principal neurons and interneurons are represented by triangles and circles, respectively. (C) Summary of AAV1-hSyn-Cre injection sites along HPC in coronal sections. Injection sites in dcHPC and vHPC are shown in cool and warm colors, respectively. (D) Micrographs showing the viral GFP expression of labeled GABAergic neurons and colocalization with the GAD67 antibody. Scale bar, 20 µm. (E) Micrographs showing the viral GFP expression and colocalization of GFP+ cells and the indicated markers. (PV+ in red, SOM+ in violet). Scale bar, 100 µm. (F) Representative schemes showing the distribution of the viral GFP-expressing interneurons in MFC receiving inputs from dcHPC (*left*, case #773) and vHPC (*right*, case #772) Cre-injections that co-express either PV or SOM across the MFC subregions. (G) Pie charts showing the proportion of GFP+ cells co-expressing PV (red) or SOM (violet) among the total GFP-labeled interneurons receiving dcHPC (*left*, n=6) or vHPC (*right*, n=4) inputs. (H-I) Subregional distribution of the GFP+ interneurons receiving HPC inputs in MFC that co-express the indicated markers PV (red) or SOM (violet) for dcHPC (H, n=6) and vHPC (I, n=4). Data are presented as violin plots, with each circle corresponding to one sample. For one sample, proportions of PV+ and SOM+ cells across the MFC subregions do not sum up to 1, with the residual proportion corresponding to unidentified GFP+ interneurons. Mann-Whitney test corrected for multiple comparisons using the Holm-Šídák method.

These results show that although dcHPC and vHPC project to different subregions in MFC, they innervate similar proportions of PV+ and SOM+ interneurons in the target subregions. However, in contrast to vHPC projections, dcHPC presented a trend to recruit a larger overall proportion of PV+ cells over other types of interneurons in MFC, which may implicate an increased perisomatic inhibition of MFC neurons by dcHPC innervation, a possibility that still needs to be tested.

## Discussion

There exists strong and extensive evidence that the direct HPC-MFC projection is critical for memory and emotion^1,2,20,24,64,65^. Most studies, however, focused on the dense projections from vHPC, and the projections from dcHPC have been largely disregarded. Our results provide a detailed and systematic documentation of the organization of the HPC projections from the entire HPC long axis, emphasizing the additional relevance of dcHPC as a main contributor to the projections to MFC. More importantly, our data point to a number of differences that would support to consider the vHPC and dcHPC components as two different parallel HPC-MFC pathways. We show that dcHPC and vHPC target different subregions in MFC: vHPC projects densely to IL, PLv and MO cortices (**Figure 7A**), whereas dcHPC targets mainly the more ventrally located DP and MO cortices (**Figure 7B**). Subsequent whole-cell recordings combined with optogenetic stimulation showed that dcHPC and vHPC innervation of projection neurons in MFC elicit excitatory and inhibitory postsynaptic responses, though with a notable difference in overall E/I balance. These observations were then confirmed and further detailed anatomically, indicating that the two pathways innervate significantly different proportions of GABAergic neurons in MFC, with dcHPC engaging a larger proportion of GABAergic cells than vHPC. In addition, we found that vHPC targets similar proportions of PV+ and SOM+ interneurons in MFC, whereas dcHPC innervates a larger proportion of PV+ interneurons.

**Figure 7.**
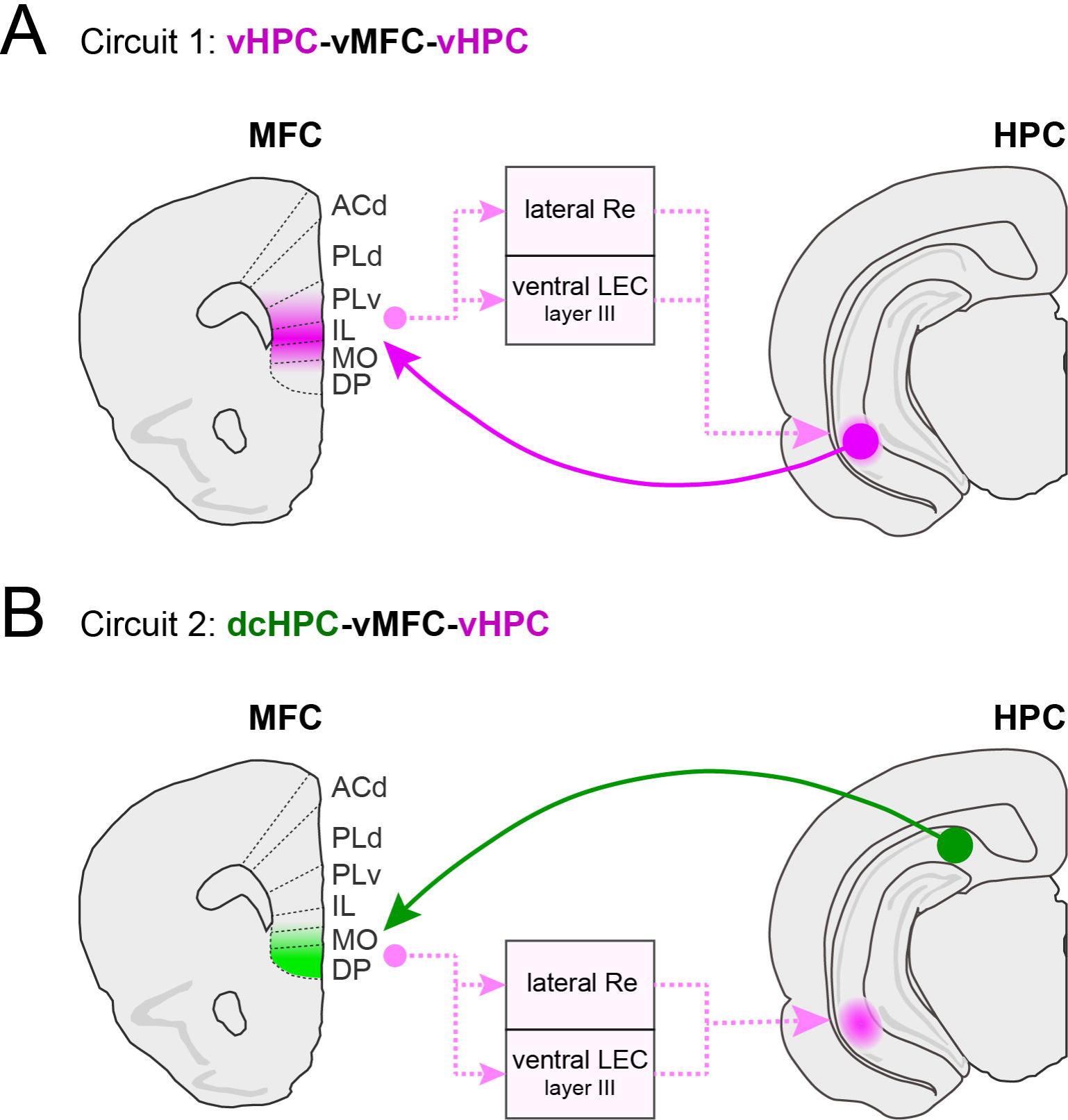
Summary of the connectivity between HPC and the subregions in MFC. (A) Reciprocal ventral circuit with a direct projection from vHPC to IL, PLv and MO in vMFC, and an indirect pathway from vMFC back to vHPC through the lateral part of Re and the ventral part of LEC layer III neurons. (B) Proposed circuit connecting dcHPC-vMFC-vHPC, with a direct projection from dcHPC to DP and MO in vMFC, and an indirect pathway from vMFC to vHPC through the lateral part of Re and the ventral part of LEC layer III neurons. *Note:* There is a reciprocal dorsal circuit by which PLd and ACd in dMFC receive limited projections from dcHPC, but sends projections back to dcHPC through the medial part of Re and the dorsal part of LEC layer III neurons.

### Organization of the direct HPC-MFC pathways along the dorsoventral axis

Direct inputs from HPC to MFC were previously described in rats as arising from restricted portions of CA1 and SUB^9,10^. These studies showed that the dorsal-rostral portion of CA1 does not project to MFC and projections originated from intermediate and ventral levels of distal CA1 and the directly adjacent proximal part of SUB, reaching AC, PL, IL and orbital areas in the frontal cortex. Although both studies reported that the density depended on the location of the injections along the longitudinal axis of HPC, it remained unclear whether there are any topographical differences in the projections from different dorsoventral HPC portions to MFC. Our anterograde tracing results show that in both mice and rats, HPC projections to MFC subregions significantly differ along the dorsoventral axis. Whereas vHPC, as previously reported^9,10^, innervates densely IL, PLv and MO subregions, dcHPC innervates the most ventral part of MFC, DP and MO cortices. Note that according to our observations, distal CA1 and proximal SUB show similar projection topographies in MFC, and therefore we analyzed the experimental data together. Corroborating this similarity between CA1 and SUB are the reported projections to MFC arising from genetically defined SUB domains in mice^66^. The latter authors observed that the dorsal-proximal part of SUB, included in dcHPC in our study, targeted mainly DP, whereas the ventral SUB projected densely to IL.

While conventional anterograde tracers allow to observe fiber distribution in target areas, the anterograde transsynaptic tracing method allows to identify the actual neurons receiving HPC inputs in MFC and visualize the projecting axons. The distribution of HPC-postsynaptic neurons across the MFC subregions replicated the fiber distributions observed in our initial anterograde tracing experiment. The dcHPC-postsynaptic neurons were mainly found in all layers of DP and MO, whereas vHPC-postsynaptic neurons were distributed across all layers of IL, PLv and MO, with a significant larger proportion in deep layers. We further observed that axon terminals from MFC projection neurons receiving HPC inputs were distributed across several subcortical areas such as the striatum, basal amygdala, mediodorsal thalamus, periaqueductal gray, and dorsal raphe, in line with previous anatomical studies^42,54,55^. Furthermore, dcHPC-receiving MFC neurons seemed to distinctively innervate the mammillary body, the ventral tegmental area, and the submedial thalamic nucleus, observations that correspond to characteristic projections previously ascribed to DP_42,67,68._

Interestingly, there is evidence that most MFC-projecting CA1 neurons have collaterals to different brain areas^69,70^, and that these long-range axon collaterals could promote synchronized neural activity and long-term synaptic plasticity in different brain areas with possible behavioral consequences^69,70^. For example, vCA1 neurons projecting simultaneously to both MFC and amygdala, induce synchronized neural activity in MFC and amygdala, and convey contextual information to basal amygdala relevant for contextual fear conditioning^69^.

### HPC activation of inhibitory circuits in MFC

Clinical studies as well as animal models suggest that alterations of cortical excitatory and inhibitory balance can give rise to social and cognitive deficits^71–73^. Several reports demonstrated that vHPC engages distinct interneuron populations in MFC to regulate the network dynamics and influence behavior ^21,25,27,74^. The present whole-cell recordings revealed that dcHPC terminal axons in MFC likewise could regulate MFC network dynamics, though with an overall more inhibitory E/I ratio in MFC than vHPC. This difference in E/I ratio could result from the fact that the dcHPC pathway innervates a larger proportion of GABAergic neurons in MFC than vHPC. Here we focused on the main classes of PV+ and SOM+ neurons, in view of their striking different laminar distribution, but VIP+ and CCK+ neurons, known to receive inputs from vHPC^57^, need to be analyzed as well.

Adding to the complexity of these inhibitory networks, reports indicate that within the hippocampal formation, the superficial and deep layers of field CA1 exhibit differences in their preferred inhibitory neuron innervations^28,57,75^. Furthermore, the differential circuits in which these interneurons in MFC are embedded appear to shape behavior differently. For example, PV+ interneurons have been implicated in avoidance behavior^76^, and more specifically vHPC innervation of PV+ interneurons in MFC facilitates formation of social memory^26^ and inhibits conditioning fear relapse^21^. In contrast, SOM+ interneurons seem to be involved in exploration and approach behavior^28,77^ and vHPC innervation of SOM+ interneurons facilitates spatial working memory^19^.

### Reciprocal circuits connecting HPC and MFC along the dorsoventral axis

In addition to the direct HPC-MFC pathways, we also identified the indirect pathways from MFC back to HPC. The retrograde transsynaptic tracing showed that these pathways disynaptically connect MFC to HPC and that they are topographically organized along the dorsoventral axis. The dorsal MFC (dMFC), including ACd and PLd, disynaptically projects to dcHPC whereas the ventral MFC (vMFC) subregions PLv and IL, disynaptically project to vHPC. These indirect pathways relay in brain regions such as the thalamic Re and layer III neurons in EC. The overall topographical organization of these disynaptic MFC-HPC pathways presented here is in line with the one described previously^39^, though we were not able to corroborate the presence of the additional relay in the anterior thalamic nuclei for the dorsal HPC pathways.

Considered together, the direct HPC-to-MFC pathways and the indirect MFC-to-HPC pathways that we describe here, consistently point to a dorsoventral organization of the bidirectional connectivity between HPC and MFC. We propose a reciprocal ventral circuit by which vMFC projects disynaptically to vHPC and receives monosynaptic inputs from vHPC (**Figure 7A**). Conversely, dMFC projects disynaptically to dcHPC, and, although direct projections from dcHPC to dMFC are very limited, dcHPC may reach dMFC disynaptically via LEC^78–80^ or thalamic Re^81,82^. The bidirectional ventral circuit (**Figure 7A**) provides routes for vMFC and vHPC to coordinate their activities and the reported shared involvement in emotional, autonomic and sympathetic functions^11,18^, including social behavior^26^, social memory^25^, context-dependent fear memory^22^, renewal of fear^21,83,84^, and anxiety-related behavior^23,24^. The dorsal circuits, connecting dMFC with dcHPC are probably involved in spatial-related cognitive functions. The dcHPC likely provides spatial information of different nature compared to proximal CA1^85,86^. Furthermore, some reports showed that dcHPC may modulate nociceptive responses in PL and AC^87^, and even strengthen fear memories through PL^35^.

Although the MFC-HPC circuits seem functionally segregated along the dorsoventral axis, i.e. a dorsal circuit related to spatial and cognitive functions and a ventral circuit related to emotion, it is likely that these circuits work together to support encoding, consolidating, and retrieving episodic memories^88,89^. MFC clearly requires some awareness of current spatial information to enable behaviors such as decision-making and flexibility. Although the interplay between the two circuits could be mediated by intrinsic connectivity within either HPC and/or MFC, our results point to an additional, unknown direct pathway, connecting dcHPC to the most ventral part of vMFC that may have a role facilitating such communication (**Figure 7B**).

### The dcHPC-vMFC circuit as a connection between dorsal-caudal and ventral HPC-MFC

In addition to the parallel ventral and dorsal circuits described above, we propose a third circuit consisting of a ‘crossed’ connectivity along the dorsoventral axis, by which dcHPC sends monosynaptic inputs to the ventral-most MFC but projects disynaptically back to vHPC (**Figure 7B**). This ventral-most MFC comprise DP and MO cortices in mice. However, whereas MO receives important innervation from both vHPC and dcHPC, DP cortex is characteristically innervated mainly by dcHPC.

Connections of DP with the rest of the brain are comparable to IL^54^. However, DP has specific connections that suggest that within the MFC subregions, DP may have a different functionality^13,90–92^. Particularly, DP projects densely to the mammillary body^67^ and receives dense inputs from LEC^93^. In addition, robust specific spatial-related information may also reach MFC through this dcHPC-DP circuit^85,94,95^. Recent studies suggest that DP plays an important role in autonomic responses and emotional-related behavior, i.e., DP seems to drive sympathetic stress responses through connections with the dorsomedial hypothalamus^14^, regulate fight-or-flight responses through its connectivity with the central amygdala^46^, and modulate affective behavior and fear memory^96,97^. The circuit through which dcHPC connects with DP may thus play an important and distinctive role in memory processing and emotional-related behavior or autonomic/sympathetic responses.

## Conclusion

We show that projections from HPC to MFC are differently organized along the dorsoventral axis. Two parallel pathways emerging from dcHPC and vHPC have been identified, which innervate distinct proportions of excitatory and inhibitory neurons across MFC. These direct projections from HPC to MFC, together with the indirect projections from MFC back to HPC, result in two circuits (**Figure 7**): a known reciprocal circuit connecting the emotional-related areas vHPC and vMFC (**Figure 7A**), and a new circuit connecting the cognitive-related dcHPC with the emotional-related vMFC, particularly DP cortex, which projects to vHPC (**Figure 7B**), thus connecting functionally different areas in HPC and MFC. The functional heterogeneity of both HPC and MFC along the dorsoventral axis suggests that the contribution of these circuits to memory processing and emotional regulation may be different. Further exploration of the functional particularities of these parallel circuits may have important implications for understanding neurological and psychiatric disorders.

## Acknowledgements

This work was supported by the following grants and organizations: Grants-in-Aid for Scientific Research on Innovative Areas (21H00178 to SO) and Grants-in-Aid for Scientific Research (23H00073 to K-IT) from the Japan Society for Promotion of Science; PRESTO (JPMJPR21S3 to SO) and Moonshot R&D (JPMJMS2292 to K-I) from the Japan Science and Technology Agency.

## Authors Contributions

PA and SO collected and analyzed the anatomical data: PA gathered data from the anterograde tracing experiments whereas SO focused on the RV retrograde tracing experiments. PA also conducted the electrophysiological experiments and performed the corresponding data analysis. All authors contributed to the conceptualization of the study and discussions that shaped the current manuscript, which was first drafted by PA and edited and reviewed by SO, MPW, and K-IT. All authors approved the final version of the manuscript.

## Supplemental information

Document S1. Figures S1-S3.

## Materials and Methods

### Animals

All animals were group housed at a 12:12 hour reversed day/night cycle and had ad libitum access to food and water. The anterograde projection patterns were characterized using adult C57BL/6N mice weighing 20-30 g (n=12 male, n=8 female), and additional adult SD rats weighing 200-390 g (n=3 male) for supplementary data. The retrograde rabies viral tracing experiments were carried out in young adult Wistar rats weighing 200-230 g (n=16 male). For the anterograde transsynaptic experiments, adult C57BL/6N mice (n=31 male) were used. The electrophysiological data was obtained from 8–10-weeks old male C57BL/6N mice (n=23 male). All animals were purchased from Japan SLC (Shizuoka, Japan). All experiments were approved by the Center for Laboratory Animal Research, Tohoku University. The experiments were conducted in accordance with the Tohoku University Guidelines for Animal Care and Use.

### Stereotaxic injections for tracer and viral transduction

Animals were anesthetized with isoflurane in an induction chamber, and were injected intraperitoneally with ketamine (80 mg/kg) and xylazine (10 mg/kg). Animals were mounted on a stereotaxic frame. Isoflurane was administered via a surgical anesthesia mask throughout the surgery at a stable level between 1-2%. Eye ointment was applied to the eyes of the animal to protect the corneas from drying out. The skin overlying the skull was disinfected with iodide, and local anesthesia was injected subcutaneously (lidocaine, 10 mg/kg). The skull was exposed, and a small burr hole was drilled above the injection site. For the anterograde tracing experiments, either 2.5% Phaseolus vulgaris-leucoagglutinin (PHA-L, Vector Laboratories L-1110) or 3.5% 10 kDa Biotinylated Dextran amine (BDA, Invitrogen-D1956) was injected by iontophoresis with 12-µA positive current pulses (6-s on, 6-s off) for 10- or 15-min using glass micropipettes, positioned differently along the longitudinal axis of HPC.

For the anterograde transsynaptic experiments, AAV1-hSyn-Cre (1.9x10^13^-2.1x10^13^ gc/ml; 200 nl; Addgene-10553) was injected in either dorsal-caudal or ventral HPC, and either AAV8-hSyn-DIO-mCherry (3.6x10^12^ gc/ml; 3x100 nl; Addgene-50459) or AAV9-hDlx-Flex-GFP (4.4x10^12^ gc/ml; 3x100 nl; Addgene-83895) was injected along the dorsoventral axis of MFC. For the electrophysiological experiments, AAV9-CaMKIIa-hChR2(H134R)-EYFP (2.2x10^12^ gc/ml; 100 nl; Addgene-26969) was injected in either dorsal-caudal or ventral HPC and for retrograde labeling AAVrg-hSyn-mCherry (2x10^13^ gc/ml; 2x100 nl; Addgene-114472) or alternatively, Red RetroBeads^TM^ IX (1:1, 2x100 nl; Lumafluor Inc.) was injected in the basal amygdala.

For retrograde transsynaptic tracing, we used recombinant rabies virus (RV) vectors based on HEP-Flury strain (Ohara et al., 2009, 2013). To examine the monosynaptic inputs to the hippocampal subfields, we used a non-transsynaptic G-deleted RV vector (150-300 nl of rHEP5.0-ΔG-mRFP (6.0×10^8^ FFU/ml), 150nl of rHEP5.0-ΔG-AcGFP1 (3.0×10^9^ FFU/ml)), or a propagation-competent RV vector (150-300 nl of rHEP5.0-CVSG-mRFP (8.0×10^8^ FFU/ml), 150-300 nl of rHEP5.0-CVSG-EGFPx2 (7.0×10^8^ FFU/ml)) with two days of survival time. Multisynaptic inputs were examined by using RV vectors with longer survival period (four to five days). Each virus was injected along with 1 % of pontamine sky blue in order to mark the injection site.

The stereotaxic coordinates relative to bregma for mice were (AP, ML, DV): dorsal-caudal HPC = -2.9, +2.1, -1.2 mm; ventral HPC = -2.9, +3, -3.2 mm; MFC = +2.1, +0.4, (-1, -1.6, -2.3) mm; BLA = -1.1, +3.2, -4.2 mm; BMA = -1.10, +2.8, -4.7 mm; and for rats were: dorsal-caudal HPC=-5.1, +4.6, =2.5 mm; ventral HPC=-5.5, +6.0, -4.6. For the viral transduction, borosilicate pipettes with 20-30 µm diameter tips were back filled with a volume of 100-200 nl of the AAV vector that was pressure-injected at 50 nl/min using a 1-µl Hamilton microsyringe. The pipette was left in place for additional 10 min, allowing time to diffuse away from the pipette tip before slowly pulling it back from the brain. The wound was sutured and after anesthesia recovery, the animal was returned to its home cage.

### Immunohistochemistry and imaging of neuroanatomical tracing samples

Survival time for injected animals was 10 days after tracer injection and 2-3 weeks after viral injections. Injected animals were deeply anesthetized with isoflurane and perfused intracardially with Ringer’s solution (0.85% NaCl, 0.025% KCl, 0.02% NaHCO_3_) followed by the fixative 4% paraformaldehyde (PFA) in 0.1 M phosphate buffer (PB) solution. The brains were then removed from the skulls, postfixed in PFA overnight at 4°C, and then cryoprotected for at least 24 h at 4°C in a 0.125 M PB solution containing 20% glycerol and 2% dimethyl sulfoxide (DMSO). The brains were cut into 40-µm thick sections in the coronal plane on a freezing microtome and collected in six equally spaced series for processing.

For immunofluorescence staining, floating sections were rinsed in phosphate buffer saline (PBS) containing 0.1% Triton X-100 (PBS-Tx), followed by a 1 h incubation in blocking solution containing 5% normal goat serum (NGS) in PBS-Tx at room temperature (RT). Sections were subsequently incubated with primary antibodies diluted in the blocking solution for 20-48 h at 4°C, washed in PBS-Tx (3x10 min), and incubated with secondary antibodies diluted in PBS-Tx for 3-5 h at RT. Finally, sections were rinsed in PBS (3x10 min), mounted on gelatin-coated slides, air-dried, cleared in xylene, and coverslipped with Entellan New (Merck Chemicals, 107961) mounting medium.

To visualize the PHA-L tracer, we used rabbit anti-PHA-L IgG (1:800, Vector Laboratories AS-2300) as primary antibody and Alexa Fluor 647 goat anti-rabbit IgG (1:400, Jackson ImmunoResearch 111-605-144) as secondary antibody. BDA was visualized with Cy3-streptavidin (1:400, Jackson ImmunoResearch 016-160-084) added along with the secondary antibodies. The primary antibodies used to visualize protein-expression were the following: rabbit anti-Cre (1:3000, Novagen 69050-3), mouse anti-Cre (1:2000, Chemicon MAB3120), rat anti-RFP (1:500, Chromotek AB_2336064), rabbit anti-DsRed (1:500, TaKaRa Bio Z2496N), mouse anti-GAD67 (1:200, Chemicon MAB5406), mouse anti-PV (1:2000, Sigma-Aldrich P3088), and rat anti-SOM (1:1000, Millipore MAB354). The respective secondary antibodies were selected among the following: Alexa Fluor 647 anti-rabbit (1:400, Jackson ImmunoResearch 111-605-144), Alexa Fluor 647 anti-mouse (1:400, Jackson ImmunoResearch 115-605-146), Cy3 anti-rat (1:400, Jackson ImmunoResearch 112-165-167), Cy3 anti-rabbit (1:400, Jackson ImmunoResearch 111-165-144), and Alexa Fluor 647 anti-rat (1:400, Jackson ImmunoResearch 112-605-003). For counterstaining, sections were stained with either guinea pig anti-NeuN (1:1000, Millipore ABN90P) or mouse anti-NeuN (1:1000, Millipore MAB377) as primary antibodies, and either DyLight 405 goat anti-guinea pig IgG (1:400, Jackson ImmunoResearch 106-475-003) or Alexa Fluor 488 goat anti-mouse IgG (1:400, Jackson ImmunoResearch 115-545-146) as secondary antibodies. Alternatively, NeuroTrace-DeepRed 640/450 (1:200, Invitrogen N21483) was used for counterstaining.

Sections were imaged using an automated scanner (Zeiss Axio Scan Z1) with a 10x/0.45 NA objective. For GAD+ cell quantification, images were scanned across a Z-stack range using confocal microscopy (Zeiss LSM-900 with Axio Observer) using a 20x/0.8 NA objective.

### Analysis of neuroanatomical tracing samples

The distribution of labeled axons or postsynaptic neurons in MFC was quantified in coronal sections spaced 240 µm apart. The mouse MFC subregions were identified based on cytoarchitectonic criteria of Werd et al.^98^. The lamination in superficial and deep layers was defined according to Gabbott et al.^99^.

For tracer labeled-fiber quantification, each MFC section was segmented into 100-µm-wide columnar bins delineated along the border between layers L3/L5. The fluorescence intensity of the labeled-fibers within each columnar bin was obtained using ImageJ. The intensity of immunolabeling in all bins was normalized to the bin with maximum intensity in the same sample. To compare the differences in projection patterns distributed across the MFC subregions, the normalized fluorescence intensities of bins were summed up within each subregion. Then, the proportion of labeled-fibers in each MFC subregion among the total observed in MFC was calculated for an injection sample, so that values from all subregions sum up to 1 for one individual sample. In figures, individual values are presented along with violin plots. Data in text are given as arithmetic mean ± SEM. The proportions in each subregion were compared using a one-way ANOVA followed by Holm-Šídák multiple comparisons test.

The proportion of Cre-labeled cells per subregion or layer was calculated as the number of Cre-labeled cells in the subregion or layer among the total of Cre-labeled cells in MFC. The proportion of PV+ or SOM+ cells per subregion was calculated as the number of PV+ or SOM+ cells in each subregion among the total of GFP-expressing cells in MFC. Data in figures are presented as individual values in violin plots, including maximum and minimum values and 25 and 75 percentile range. Data in text are presented as median. The values were compared using Mann-Whitney test corrected for multiple comparisons using the Holm-Šídák method.

The proportion of either GAD+ or GAD-cells per subregion was calculated as the number of GAD+ or GAD-cells in each subregion among the total of mCherry-expressing neurons in MFC. The proportions were compared in each subregion for an injection case using a two-tailed t-test.

All statistical analysis were carried out using GraphPad Prism 10. Thresholds for significance were set at *p<0.05, **p<0.01, ***p<0.001, and ****p<0.0001.

### Slice preparation

After 2-3 weeks of viral expression, mice were deeply anesthetized with isoflurane before being sacrificed. After decapitation, brains were rapidly removed and placed in an ice-cold oxygenated cutting solution containing (in mM): 110 Choline-Cl, 2.5 KCl, 7 MgCl_2_, 0.5 CaCl_2_, 25 Glucose, 1.25 NaH_2_PO_4_, 25 NaHCO_3_, 11.5 Na-ascorbate, 3 Na-pyruvate, 100 D-Mannitol. Coronal slices of MFC (300-µm thick) were cut on a vibratome (DSK Linearslicer PRO 7) keeping the brain immersed in ice-cold cutting solution. Then, slices were transferred to artificial cerebrospinal fluid (ACSF) containing (in mM): 126 NaCl, 3 KCl, 1.2 NaH_2_PO_4_, 10 Glucose, 26 NaHCO_3_, 3 MgCl_2_, 0.5 CaCl_2_, bubbled with O_2_. Slices were kept for 30 min at 35°C, before being allowed to recover for at least 30 min at room temperature. All chemicals were purchased from either Sigma-Aldrich or Wako.

### Electrophysiology

Recordings were conducted at 28-30°C in oxygenated ACSF containing (in mM): 126 NaCl, 3 KCl, 1.2 NaH_2_PO_4_, 10 Glucose, 26 NaHCO_3_, 1.5 MgCl_2_, 1.6 CaCl_2_ (∼295 mOsm). For each individual slice, pyramidal neurons located in MFC subregions were identified by infrared-differential interference contrast (IR-DIC) microscopy. Amygdala-projecting neurons were selected for recording by the mCherry-expression, result from retrograde AAV transduction, observed under fluorescent illumination. Whole-cell voltage-clamp recordings were performed using borosilicate glass pipettes (2-4 MΩ) filled with a Cs-based intracellular solution containing (in mM): 120 Cs-gluconate (Hellobio, HB4822), 10 HEPES, 10 Na-phosphocreatine, 4 Mg-ATP, 0.4 Na-GTP, 0.5 EGTA, 2 QX314, 10 TEA-chloride (pH ∼7.3, CsOH; ∼290 mOsm). Biocytin (2.5%) was included in the internal solution, which was allowed to diffuse throughout the cell for posterior confirmation of location and cell morphology by immunolabeling. Electrophysiological recordings were made with a Double Integrated Patch Amplifier (IPA) and SutterPatch software (Sutter Instrument), filtered at 2 kHz and sampled at 10 kHz. The initial series resistance was <20 ΩM, and recordings ended if series resistance rose >25 ΩM. Excitatory and inhibitory postsynaptic currents (EPSC and IPSC) were recorded at -70 mV and +10 mV, respectively. In some experiments, to isolate the monosynaptic inputs, TTX (1 µM), 4-AP (100 µM), and elevated Ca^2+^ (4 mM) were included in the recording solution, a mix that blocks the action potentials but restores presynaptic release^100,101^. In other experiments, the excitatory responses were isolated by including gabazine (10 µM) in the recording solution to block the GABA-_A_ receptors. All chemicals were purchased from either Sigma-Aldrich or Wako.

### Optogenetic stimulation

Channelrhodopsin-2 (ChR2) was expressed in presynaptic neurons and their terminal axons in MFC were activated with a brief light pulse from a blue LED (473 nm, Zeiss Colibri 7). For wide-field illumination, light was transmitted via a 40x/0.8 NA objective. For each experiment, LED pulse power and duration were adjusted to a typical value of 15 mW and 2 ms, respectively.

### Immunohistochemistry and imaging of recorded slices

Slices containing biocytin-filled cells were fixed in 4% PFA overnight at 4°C. For staining, slices were washed with PBS-Tx (0.3%; 5x15 min), then incubated in 10% NGS blocking solution in PBS-Tx for 3 h at RT, and then incubated in the same blocking solution containing a diluted mixture of primary antibodies at 4°C for 4 days. Subsequently, slices were washed with PBS-Tx (0.3%; 5x15 min) and incubated in a dilution of secondary antibodies diluted in PBS-Tx overnight at RT. Slices were then washed in PBS-Tx (3x15 min) and stored in PBS at 4°C. Slices were dehydrated by subsequent 10-min exposure to ethanol/distilled water mixtures of 30%, 50%, 70%, 90%, then two times of 100% ethanol, and a (1:1)-mixture of ethanol/methyl salicylate. Slices were then cleared with methyl salicylate and embedded on metal slides with coverglass. Dehydration and clarification were not applied when using RetroBeads as a tracer. Z-stack images were collected with confocal microscope (Zeiss LSM-900 with Axio Observer) using a 20x/0.8 NA objective.

GFP-expression was enhanced with the primary antibody chicken anti-GFP (1:500, Abcam AB13970) and secondary antibody Alexa Fluor 488 anti-chicken (1:500, Invitrogen A-11039). mCherry-expression was enhanced using primary antibody rabbit anti-DsRed (1:300, TaKaRa Bio Z2496N) and secondary antibody Cy3 anti-rabbit (1:200, Jackson ImmunoResearch 111-165-144). Biocytin was visualized with Alexa Fluor 647 streptavidin (1:200, Jackson ImmunoResearch 016-600-084) added to secondary antibody mix. For counterstaining, the primary antibody used was guinea pig anti-NeuN (1:500, Millipore ABN90) and secondary antibody DyLight 405 anti-guinea pig (1:200, Jackson ImmunoResearch).

### Analysis of electrophysiological data

EPSC and IPSC events were analyzed off-line using the Synaptic Event Analysis module from the SutterPatch application (Igor Pro software). Quantitative data from multiple slices are given as median and 25^th^, 75^th^ percentile. Data in figures are presented as individual values indicating median in violin plots showing minimum and maximum values, with 25-75 percentile range. Data in text are only given as medians. Statistical analysis was performed using GraphPad Prism 10 software. Mann-Whitney test was used and corrected for multiple comparisons using the Holm-Šídák method. Thresholds for significance were set at *p<0.05, **p<0.01, and ***p<0.001.

