## Supplemental Figures for "Dorsal-caudal and ventral hippocampus target different cell populations in the medial frontal cortex in rodents"

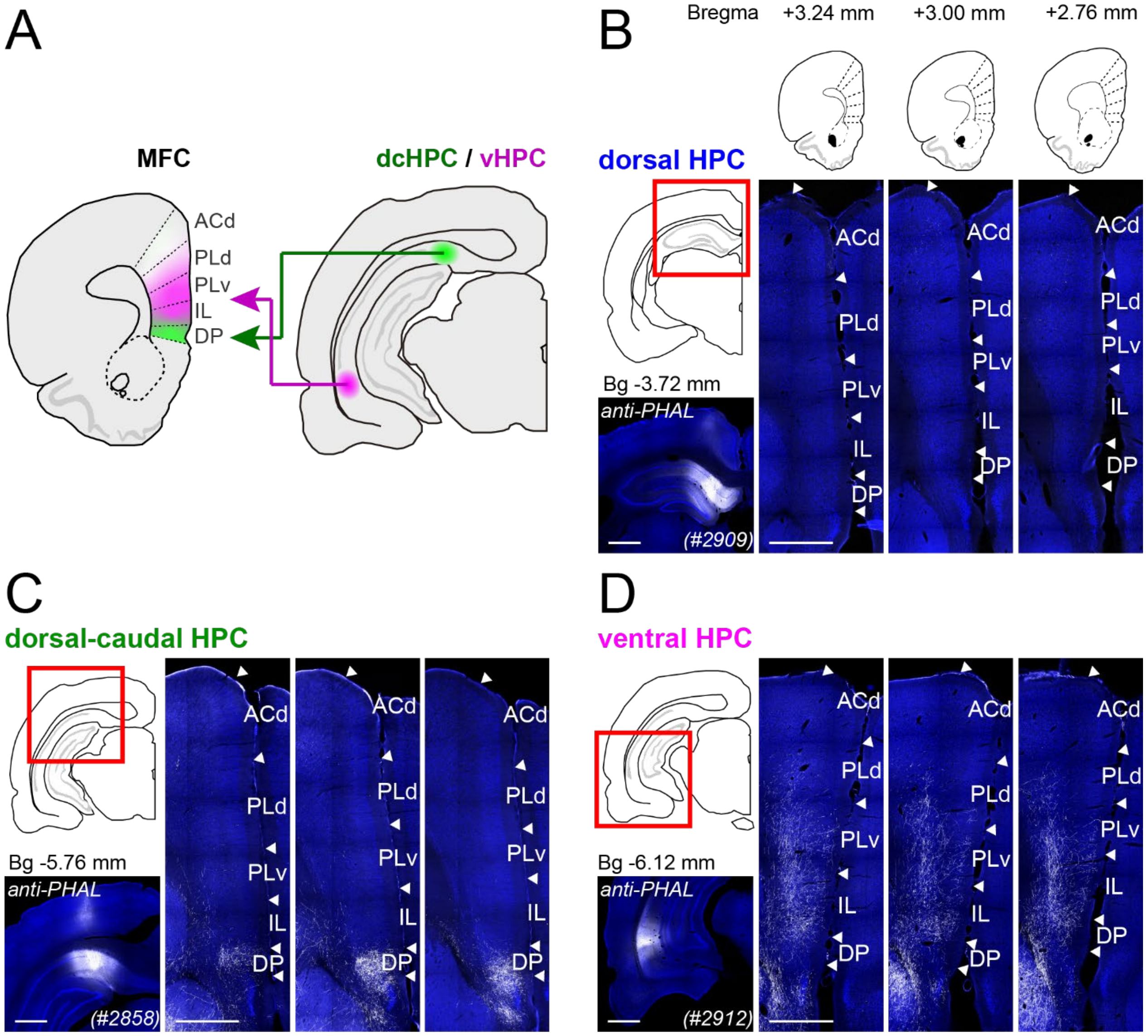


**Figure S1. Fibers from anterograde tracer injections along the dorsoventral axis of HPC are differently distributed across MFC subregions in rats.**

(A) Summary illustration of parallel pathways from dorsal-caudal and ventral HPC to MFC subregions. (B-D) Representative samples of anterogradely labeled fibers in MFC (at three coronal levels as shown in top inset) for injections in the most dorsal HPC (#2909-PHAL, B), dorsal-caudal HPC (#2858-PHAL, C), and ventral HPC (#2912-PHAL, D). Scale bars, 1 mm.

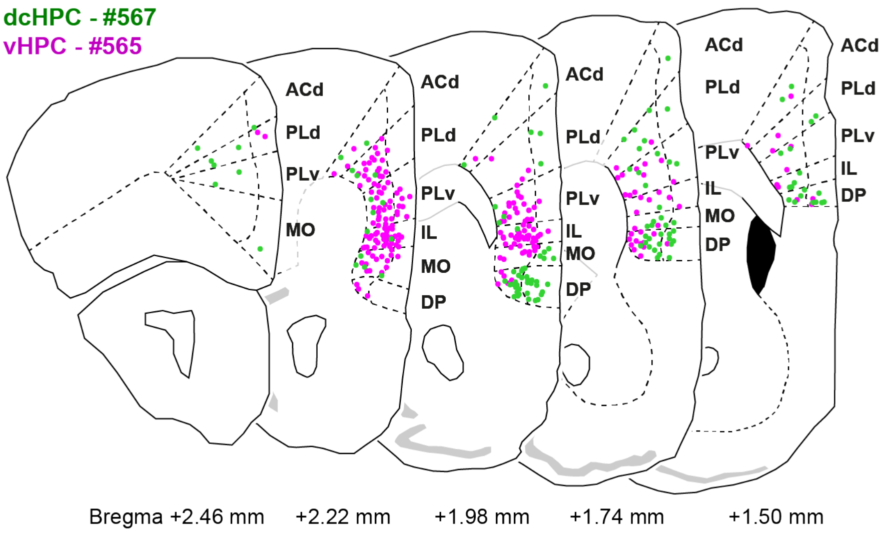


**Figure S2. Distribution of postsynaptic Cre-labeled cells throughout MFC in mice.**

Representative scheme showing the distribution of postsynaptic Cre-labeled neurons across the anterior-posterior axis of MFC for one dcHPC (case #567, green) and one vHPC (case #565, magenta) injection.

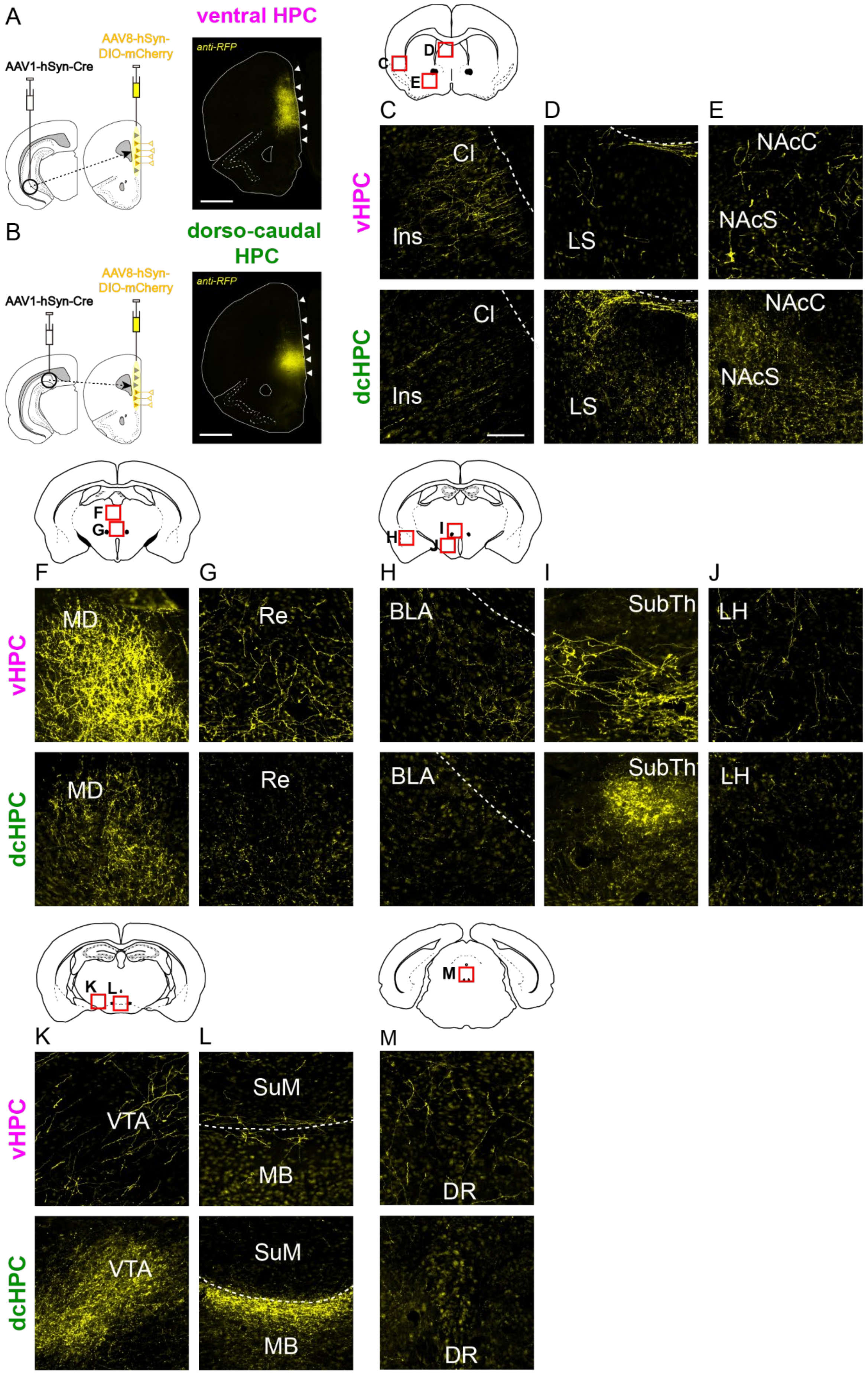

A
AAV1-hSyn-Cre
AAV8-hSyn-DIO-mCherry
ventral HPC
anti-RFP
B
dorso-caudal HPC
C
D
E
Cl
Ins
LS
NAcC
NAcS
vHPC
dcHPC
F
G
H
I
J
MD
Re
BLA
SubTh
LH
K
L
M
VTA
SuM
MB
DR

**Figure S3. Brain regions targeted by MFC neurons receiving inputs from dorsal-caudal or ventral HPC in mice.**

(A-B) Representative image showing mCherry-labeled neurons in MFC receiving inputs from ventral HPC (vHPC, A) or dorsal-caudal HPC (dcHPC, B). Note that vHPC postsynaptic neurons are mainly distributed across MO, IL and PLv subregions whereas dcHPC postsynaptic neurons are mainly distributed across DP and MO subregions in MFC (white arrow heads indicate the borders between subdivisions). (C-M) Presence of mCherry+ terminals from MFC neurons innervated by vHPC (top) or by dcHPC (bottom) in claustrum (Cl) and insular cortex (Ins, C), lateral septum (LS, D), nucleus accumbens core (NAcC) and shell (NAcS, E), mediodorsal thalamus (MD, F), nucleus reuniens (Re, G), basolateral amygdala (BLA, H), submedial thalamic nucleus (SubTh, I), lateral hypothalamus (LH, J), ventral tegmental area (VTA, K), mammillary and supramammillary body (MB, SuM, L), and dorsal raphe (DR, M). Scale bars in (A-B), 1000 μm; in (C-M), 100 μm.
